# Reference-Informed Spatial Domain Detection Using Weak Supervision for Spatial Transcriptomics

**DOI:** 10.1101/2025.09.11.675689

**Authors:** Xin Ma, Weijia Jin, Qing Lu, Ramon C. Sun, Li Chen

## Abstract

One of the key objectives in spatial transcriptomics (ST) studies is to map the complex organization and functions of tissues. We introduce GraphScrDom, a reference-informed and weakly supervised contrastive learning model that uniquely integrates expert-provided manual annotations (i.e., scribbles) on spatial grids or histology images with cell type–specific gene expression profiles derived from reference single-cell RNA-seq data to perform tissue segmentation. With only limited scribble annotations, GraphScrDom consistently outperforms existing methods across various ST platforms and at both bulk and single-cell resolutions, as evaluated by six widely used metrics, demonstrating strong generalizability and robustness. Additionally, we have developed an integrative software toolkit that includes an interactive annotation interface and a model training module for spatial domain detection, providing a unified and user-friendly framework to facilitate spatial domain analysis.

## Introduction

Spatial transcriptomics (ST) is a fast-evolving technology that enables the measurement of the gene expression while preserving the spatial context within a tissue section. In contrast to single-cell RNA-seq data (scRNA-seq), ST provides an additional spatial dimension, capturing gene expression at the level of individual cells or small groups of cells (spots) within two- or three-dimensional tissue architecture. This added spatial context offers powerful insights into studying tissue organization, developmental processes, and disease microenvironments *in situ* (Cui et al., 2025; Janesick et al., 2023; Rao et al., 2021; Sampath Kumar et al., 2023). ST technologies are generally categorized into two main approaches: sequencing and imaging-based approaches. Sequencing-based methods include next-generation sequencing platforms such as 10x Visium (10x Genomics, 2025), Slide-seq (Rodriques et al., 2019) and Stereo-seq (Chen et al., 2022), as well as in situ sequencing such as STARmap PLUS (Shi et al., 2023). Imaging-based platforms include MERFISH (Chen et al., 2015), seqFISH (Shah et al., 2016) and STARmap (Wang et al., 2018). These technologies vary in terms of gene panel coverage, ranging from hundreds to thousands of genes, and spatial resolution, spanning spot-, single cell- and subcellular level.

Identifying spatial organization of complex tissues, such as anatomical regions in the brain, tumor microenvironments in cancer, and layered structures in developing embryos, is a central goal of spatial transcriptomics (ST) studies (Jin et al., 2024; Maynard et al., 2021; Srivatsan et al., 2021). Spatial domain detection enables researchers to delineate anatomical boundaries, uncover domain-specific transcriptional programs, and better understand cellular function and identity within the spatial context. However, accurate spatial domain detection remains a challenging task, and ongoing efforts continue to explore improved methodologies. Several approaches have been developed for this purpose. The Leiden algorithm, implemented in the SCANPY package(Wolf et al., 2018), is a community detection method originally developed for single-cell clustering but also widely used for spatial domain identification (Lin et al., 2024). BayesSpace adopts a fully Bayesian model that incorporates spatial neighborhood information for refining clustering outcomes (Zhao et al., 2021). More recently, deep learning-based methods have been proposed to learn the spot- or cell-level embedding that are subsequently clustered into spatial domains. For example, SpaGCN integrates gene expression, spatial coordinates and histology image features through a graph convolutional network to derive the node embedding (Hu et al., 2021). STAGATE employs a graph attention autoencoder to learn low-dimensional latent embeddings by jointly modeling spatial proximity and gene expression profiles (Dong & Zhang, 2022). GraphST introduces a self-supervised contrastive learning within graph neural network to enhance the feature representation for spatial domain detection (Long et al., 2023).

While effective in many settings, most unsupervised approaches rely solely on spot-level gene expression, often neglecting two valuable external sources of information-histology images and cell type compositions that can potentially enhance performance. Histology images, such as H&E-stained sections, are typically generated alongside spatial transcriptomics (ST) data (e.g., 10x Visium) or can be profiled separately if unavailable. These images offer critical morphological context, enabling interpretation of gene expression in relation to tissue structure, cell types and pathology. For example, in cancer studies, histology images help distinguish tumor core, invasive margin, and stromal regions, each exhibiting distinct molecular signatures. When aligned with spatial gene expression profiles on the same tissue section, histology image provides an essential spatial reference that facilitates mapping transcriptomic patterns to anatomical features. Despite the potential, histology images are rarely leveraged explicitly in spatial domain detection, with few exceptions such as using them to refine spot-embeddings (Hu et al., 2021). Similarly, most existing methods overlook cell type composition, even though it is more biologically relevant to tissue architecture than transcriptomic heterogeneity alone.

Recent studies have demonstrated that cell type abundance maps can effectively delineate spatial structures such as cortical layers from mouse brain, using both single-cell resolution platforms (e.g., smFISH, MERFISH) and spot-resolution platforms (e.g., 10x Visium) (Biancalani et al., 2021; Kleshchevnikov et al., 2022). This is further supported by IRIS (Ma & Zhou, 2024), which improve the spatial domain detection by integrating deconvolved cell type composition (Gaspard-Boulinc et al., 2025; Li et al., 2023). The availability of rich, tissue-specific single-cell RNA sequencing (scRNA-seq), from public resources, including Human Cell Atlas (Regev et al., 2017), Tabula Muris (Tabula Muris et al., 2018), Genotype-Tissue Expression (GTEx) project (Eraslan et al., 2022), and Gene Expression Omnibus (GEO), has made this integration increasingly feasible. Another important yet underutilized resource is histology-based annotation. While polygonal annotations of anatomical structures by pathologists are standard practice, they are time-consuming, labor-intensive, and error-prone, especially in heterogeneous tissues with complex morphology (Zhang et al., 2023). A practical alternative is scribble annotation, which involves drawing freeform strokes over limited regions of interest. Scribble-based weakly supervised learning—widely adopted in medical image segmentation (Li et al., 2024; Qu et al., 2024) and histology image segmentation (Oh et al., 2025), combines these sparse annotations with semi-supervised learning to infer full-image labels. In the context of ST, scribble annotations allow pathologists to label a small subset of spots or cells efficiently, and a learning algorithm can propagate these labels across the tissue section. A recent method, ScribbleDom (Rahman et al., 2023) exemplifies this approach by leveraging sparse scribble annotations and deep learning to perform robust spatial domain detection in ST datasets.

Despite the demonstrated success of ScribbleDom and IRIS, both approaches face important limitations. IRIS depends heavily on the availability and quality of reference scRNA-seq data, and its performance may be compromised when the reference is poorly matched to the ST dataset. ScribbleDom, on the other hand, employs a convolutional neural network (CNN) classifier that requires the non-rectangular tissue section to be padded into a regular grid, potentially introducing artifacts and reducing model fidelity. This architectural constraint restricts ScribbleDom’s applicability to grid-based spatial platforms (e.g., 10x Visium), rendering it incompatible with non-grid platforms such as MERFISH. Furthermore, there is a lack of an integrated and user-friendly computational pipeline that enables human experts (e.g., pathologists) to perform both scribble annotations and spatial domain detection within a unified framework. To address these limitations, we present GraphScrDom, a unified and versatile deep learning framework and software toolkit for spatial domain detection in ST studies. GraphScrDom integrates gene expression, cell type composition, and spatial coordinates under a scribble-driven weakly supervised learning paradigm. It offers five key innovations. First, GraphScrDom constructs spatial graphs based on spot or cell coordinates without assuming a regular grid layout (e.g., hexagonal grid for 10x Visium) or rectangular tissue shape. This makes it broadly applicable to diverse ST platforms, including both grid-based and non-grid-based technologies. Second, it provides a principled mechanism to jointly model transcriptomic heterogeneity and cell type composition. Cell type compositions are inferred from reference scRNA-seq data via deconvolution, and a gated fusion module is introduced to adaptively balance the contributions of both data sources. This allows GraphScrDom to remain robust even when the reference scRNA-seq differs from the ST data due to batch effects or technical noise. Third, the framework incorporates a computer vision technique known as scribble annotation within a graph neural network–based weakly supervised learning model. Scribbles are sparse, freeform strokes drawn over regions of interest that annotate only a small portion of the tissue image. In ST applications, pathologists can apply scribbles directly onto histology images, quickly highlighting tissue regions of interest without the need for exhaustive polygonal contours. This annotation strategy is not only efficient but also reduces expert burden and mitigates potential boundary labeling errors, making it highly practical for spatial omics and histopathology. Fourth, unlike traditional two-step methods that decouple representation learning from clustering for identifying spatial domains, GraphScrDom performs both tasks simultaneously. By integrating representation learning and spatial domain inference into a single training process, the model learns embeddings that are inherently task-relevant and spatially coherent. Finally, GraphScrDom includes an end-to-end, user-friendly software toolkit that supports interactive scribble annotation and spatial domain detection in one seamless workflow, making it accessible and practical for biomedical researchers and clinical experts alike. As a result, GraphScrDom demonstrates superior performance across diverse tissues and six publicly available ST datasets, comprising 25 tissue sections from human and mouse samples profiled using five different platforms (10x Visium, MERFISH, STARmap, osmFISH, and BaristaSeq). Its combination of high accuracy, broad applicability, and interactive software support establishes GraphScrDom as a powerful and indispensable tool for tissue segmentation and spatial domain analysis in ST research.

## Results

### Overview of GraphScrDom

Existing spatial domain detection methods primarily rely on transcriptomic heterogeneity to delineate regions characterized by internal expression homogeneity and distinct boundaries from neighboring areas. However, cell type compositional heterogeneity—though closely related to transcriptomic variation—offers a more interpretable and biologically grounded view of tissue organization and remains underutilized in spatial domain segmentation. To illustrate the complementary strengths of transcriptomic profiles and cell type compositions, we performed an exploratory analysis on a representative tissue slice (#151673) from the Human Dorsolateral Prefrontal Cortex (DLPFC) ST dataset. We first examined the spatial expression patterns of several spatially variable genes to assess their utility in demarcating anatomical regions within the slice. As shown in **Figure 1A**, CNP expression is highly specific to the white matter (WM), and PCP4 is predominantly expressed in cortical layer 5. Other genes, such as NEFM, HOPX, and HPCAL1, exhibit enriched expression in multiple cortical layers: NEFM in layers 3 and 4, HOPX in layers 2 and 3, and HPCAL1 in layers 2 and 6. These patterns highlight the informative yet sometimes diffuse nature of transcriptomic signals for anatomical delineation. In contrast, the spatial distribution of five representative cell types inferred from reference scRNA-seq data shows even more distinct and localized enrichment patterns (**Figure 1B**). Specifically, Oligo_02 is exclusively localized in the WM, while Excit_11 and Excit_09 are enriched in layer 5, Excit_02 in layer 3, and Excit_08 in layer 2. These results demonstrate that cell type composition can serve as a more precise marker of anatomical structure than transcriptomic expression alone. Importantly, the clear region-specific enrichment of cell types underscores their complementary value in spatial domain detection and supports the integration of both transcriptomic and cellular composition information for improved segmentation of complex tissues.

**Figure 1.**
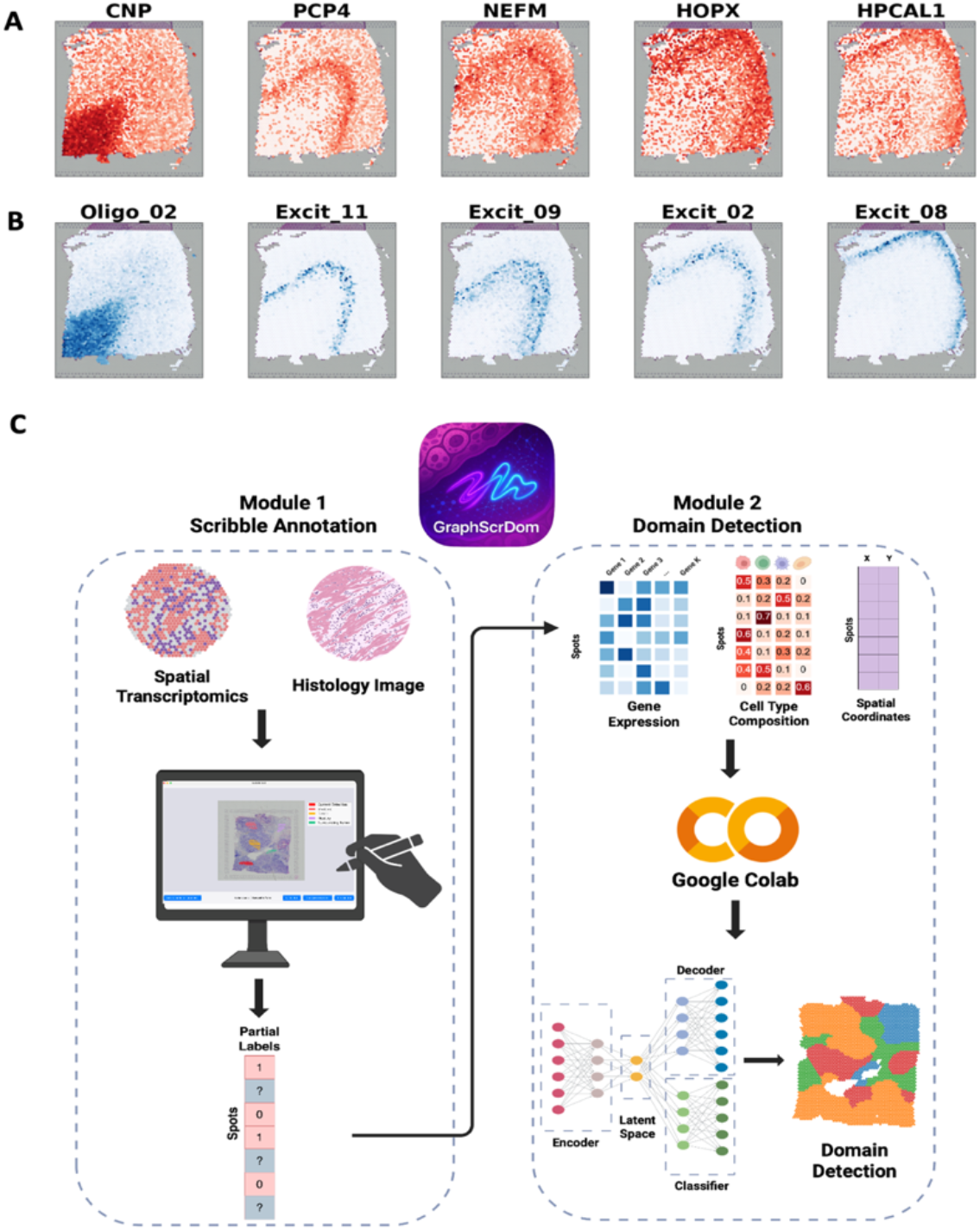
**A**. Spatial gene expression of five genes, including CNP, PCP4, NEFM, HOPX, HPCAL1, for slice #151673 in Human DLPFC spatial transcriptomics (ST) dataset. **B**. Spatial distribution of five cell types, including Oligo_02, Excit_11, Excit_09, Excit_02, Excit_08, inferred from reference single-cell RNA-seq dataset. **C**. Schematic of the GraphScrDom framework. Human experts (e.g., pathologists) perform scribbles annotations based on histology images using the integrated scribble annotation tool provided by GraphScrDom. The model takes as input the transcriptomic profiles from ST data, cell type compositions derived from reference scRNA-seq, and scribble annotations. Model training is performed via the interactive module hosted on Google Colab provided by GraphScrDom. The output is the predicted spatial domain labels for all spots on the tissue section.

Motivated by these observations, we developed GraphScrDom, a unified graph-based deep learning framework that simultaneously learns spatially informed latent representations and performs semi-supervised classification in an end-to-end manner (**Figure 2A**). GraphScrDom integrates prior knowledge from both scribble annotations and single-cell reference data within a graph neural network to enhance spatial domain detection. Specifically, both the transcriptomic profile and the inferred cell type composition are used as inputs to two modality-specific encoders—each implemented as a Self-supervised Contrastive Learning Graph Convolutional Network (SCL-GCN)—to learn latent representations that capture spatially coherent biological variation. Each SCL-GCN encoder independently learns a modality-specific representation, which is reconstructed through its corresponding decoder to preserve modality fidelity (i.e., transcriptomic profile or cell type composition). Within each encoder, a contrastive learning mechanism further refines the representations by maximizing agreement between features of neighboring nodes, thereby enhancing local structural coherence in the latent space (**Figure 2B**). The two modality-specific embeddings are subsequently integrated using a gated fusion mechanism, yielding a unified representation that balances contributions from both modalities. This fused representation is then fed into two downstream components: a decoder that reconstructs the transcriptomic profile and a semi-supervised classifier that predicts spatial domain labels. The classifier is guided by scribble annotations, enabling efficient and weakly supervised learning of spatial domains. A detailed description of the model architecture and training procedure is provided in the Methods section.

**Figure 2.**
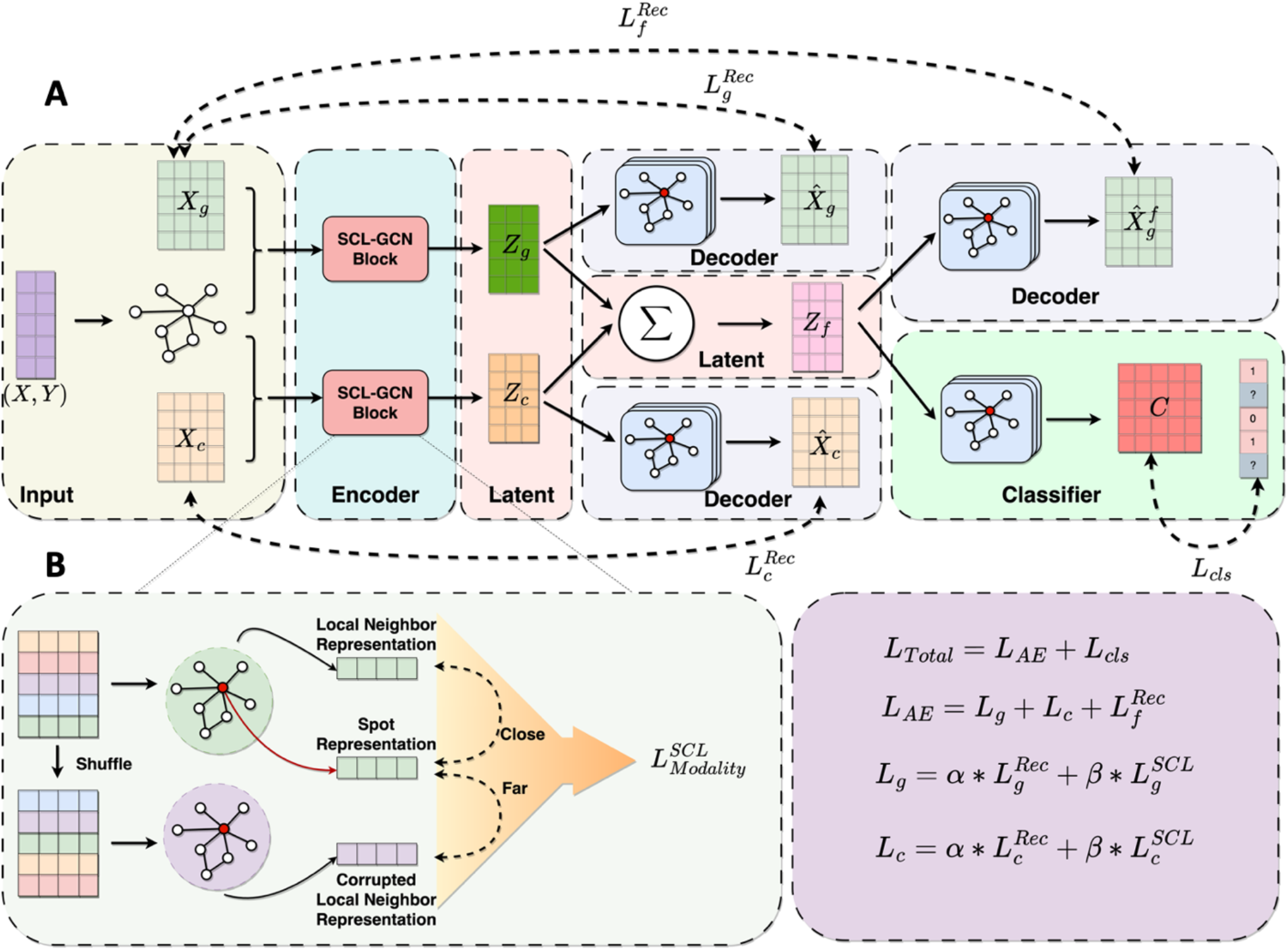
Overview of GraphScrDom. A. The model takes spatial gene expression *X*_*g*_, cell type composition *X*_*c*_, and spatial coordinates (*X, Y*), as input. Spatial coordinates are used to generate a shared graph for GCN. Each modality is encoded using a Self-contrastive Learning Graph Convolutional Network (SCL-GCN) block to produce latent embeddings *Z*_*g*_, and *Z*_*c*_. These embeddings are used for three parallel tasks: (1) modality-specific reconstruction via decoders to reconstruct 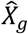 and 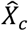, (2) gated fusion into a combined representation *Z*_*f*_, followed by reconstruction into 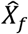, and (3) domain classification using the fused representation, guided by scribble-based supervision. B. Illustration of the Self-contrastive Learning (SCL) module used within each SCL-GCN block. For each spot, a local neighbor representation is constructed from spatially adjacent spots. A corrupted neighbor representation is created by shuffling features, and the model learns to bring positive pairs closer while pushing apart negative pairs in the latent space.

### Simulation study for multimodal integration

To demonstrate the advantage of integrating both spot-level transcriptomic profiles and cell type compositions for spatial domain detection, we designed a simulation strategy that considers varying levels of signal-to-noise ratio (SNR) for each modality. This simulation framework is intended to assess whether model performance improves when both modalities are informative, and whether the model remains robust when one modality is noisy. Specifically, we simulate a tissue slice composed of three spatial domains, whose spatial layout is derived from sample #151507 of the human DLPFC dataset. Approximately 5% of spots within each domain are assigned scribble annotations, positioned near the center of each domain to reflect typical manual labeling. Each domain contains on average ∼1,400 spots. Domain-specific transcriptomic profiles are generated using sets of spatially variable genes (SVGs) that are uniquely expressed in each domain. For each spot, the transcriptomic profile is defined as the aggregate expression of the SVGs associated with its corresponding domain. To control SNR, Gaussian noise with mean 0 and standard deviation (σ) of either 3 (low noise) or 25 (high noise) is added across all spots. After noise addition, values are clipped to ensure non-negativity. For cell type composition, we assume five cell types per spot and simulate domain-specific compositions using Dirichlet distributions. Without loss of generality, we define domain enrichment as follows: Domain 1 is enriched for cell types 1, 2, and 3; Domain 2 for cell types 2, 3, and 4; and Domain 3 for cell types 3, 4, and 5. Cell type compositions for each spot are sampled from domain-specific Dirichlet distributions: Dir(10, 5, 5, 1, 1) for Domain 1, Dir(1, 5, 10, 5, 1) for Domain 2, and Dir(1, 1, 5, 5, 10) for Domain 3. To introduce variability in cell type composition, we apply Gaussian noise with mean 0 and standard deviation of either 0.3 (low noise) or 1 (high noise), followed by renormalization to ensure the compositions sum to one. We define three simulation scenarios to evaluate model robustness: (i) both transcriptomic profile and cell type composition have high SNR; (ii) transcriptomic profile has low SNR, while cell type composition has high SNR; (iii) transcriptomic profile has high SNR, while cell type composition has low SNR. To perform an ablation study and assess the contribution of each modality, we construct two variants of GraphScrDom: **GraphScrDom_GE**, which uses only the transcriptomic profile, and **GraphScrDom_CT**, which uses only the cell type composition. These experiments allow us to quantify the effectiveness and robustness of multimodal integration in spatial domain detection.

To clearly illustrate the SNR of the two modalities in our simulation, we visualized both the transcriptomic profiles and cell type compositions across the three spatial domains using two heatmaps: one representing spot-by-gene expression for SVGs and the other representing spot-by-cell type compositions. When both modalities are of high SNR, the domain-specific transcriptomic profiles and cell type compositions align strongly along the diagonal of their respective heatmaps, indicating clear domain separation (**Figure 3A**). We then compared the predicted spatial domain maps produced by the three model variations—GraphScrDom, GraphScrDom_GE (transcriptomic profile only), and GraphScrDom_CT (cell type composition only)—along with the ground truth and scribble annotations (**Figure 3B**). Additionally, model performance was evaluated using six benchmark metrics (**Figure 3C**). Under this scenario, where both modalities are informative, all three models achieved high accuracy, with GraphScrDom attaining an ARI of 0.975, followed by GraphScrDom_CT at 0.906 and GraphScrDom_GE at 0.833. Similar trends were observed across the other evaluation metrics (**Figure 3C**). These results underscore the effectiveness of scribble-based weak supervision; with fewer than 5% of spots labeled, the models could still accurately predict spatial domains. The outcome aligns with expectations from the simulation design: SVGs expressed uniquely within domains enable reliable segmentation, and domain-specific cell type compositions also provide strong discriminatory power. Notably, GraphScrDom outperformed both single-modality variants, demonstrating that combining both modalities yields additional performance gains when both are informative.

**Figure 3.**
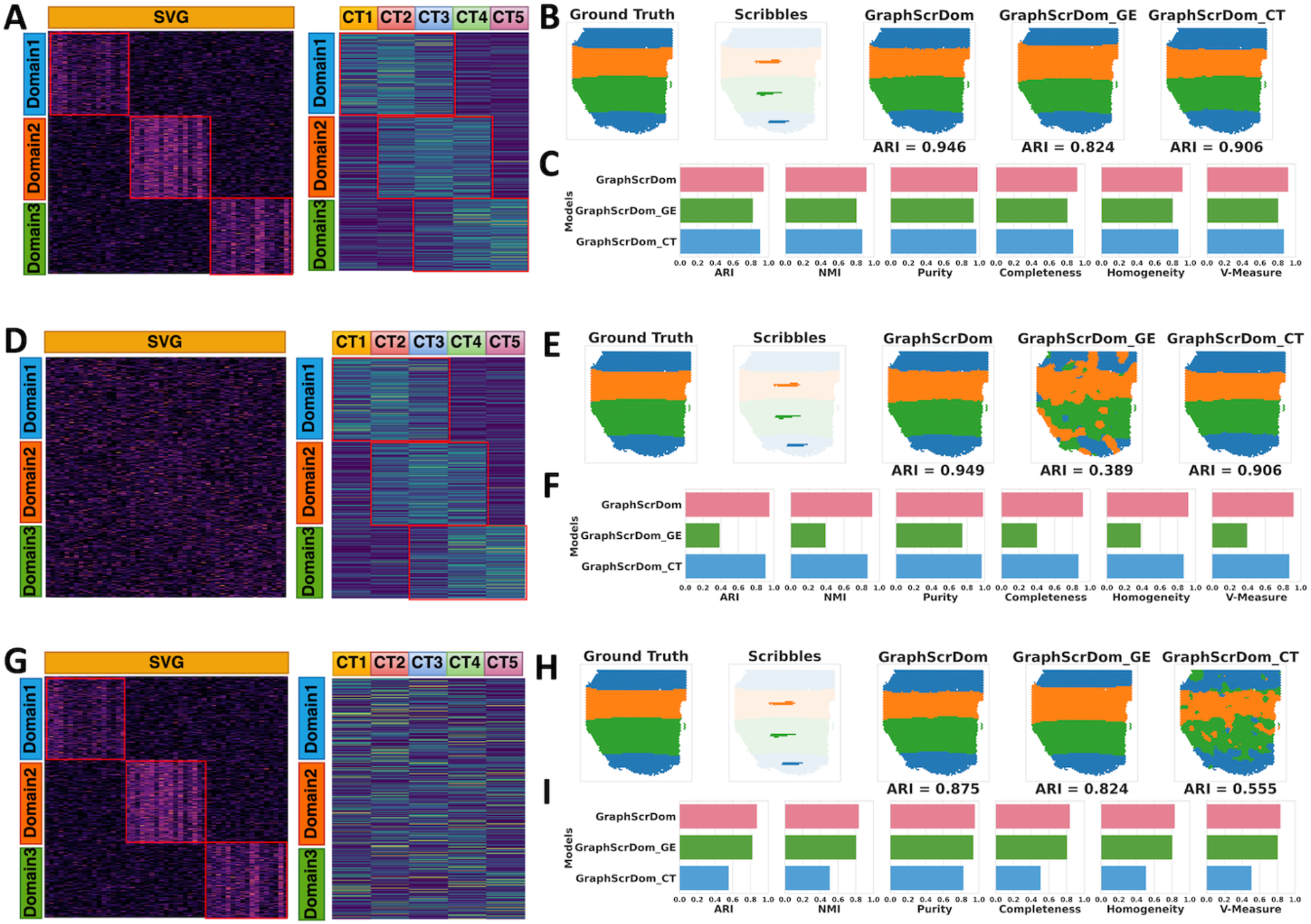
Simulation study demonstrates the effectiveness and robustness of multimodal integration in GraphScrDom. **A**. Heatmaps for Scenario i, which show simulated spatially variable gene expression (left) and cell type composition (right) under high signal-to-noise ratio (SNR) conditions across three spatial domains. **B**. Spatial maps displaying ground truth, scribble annotations, and predicted domains from GraphScrDom (multimodal), GraphScrDom_GE (transcriptomic profile only), and GraphScrDom_CT (cell type composition only) for Scenario i. **C**. Performance metrics including Adjusted Rand Index (ARI), Normalized Mutual Information (NMI), Completeness, Homogeneity, V-Measure, and Accuracy across the three model variants for Scenario i. **D**. Heatmaps for Scenario ii, where gene expression is corrupted by high noise (left), and cell type composition remains high SNR (right). **E**. Spatial domain predictions in Scenario ii across the three models. **F**. Performance metrics across the three model variants for Scenario ii. **G**. Heatmaps for Scenario iii, where gene expression remains high SNR (left), and cell type composition is corrupted by noise (right). **H**. Spatial domain predictions in Scenario iii across the three models. **I**. Performance metrics across the three model variants for Scenario iii.

We further evaluated robustness in Scenario ii, where the transcriptomic profile was degraded by high noise while cell type composition remained high SNR. As expected, no clear domain-specific pattern was observed in the SVG expression heatmap, while the domain-specific structure was preserved in the cell type composition (**Figure 3D**). Under this setting, GraphScrDom_GE’s performance dropped substantially (ARI = 0.389), with evident misclassifications between domains 1 and 3 (**Figure 3E**). In contrast, GraphScrDom achieved an ARI of 0.949—comparable to its performance in Scenario i—demonstrating strong resilience to noisy transcriptomic input. Across all six evaluation metrics, GraphScrDom consistently outperformed the other variants (**Figure 3F**), confirming its robustness and ability to denoise transcriptomic noise by effectively integrating cell type information. While GraphScrDom_CT performed reasonably well due to the clean cell type composition, it lacked the fine-grained transcriptomic variation available in GraphScrDom, resulting in slightly reduced performance. This robustness can be attributed to GraphScrDom’s adaptive gated fusion mechanism, which dynamically weighs the contribution of each modality. The learned gating function enables the model to upweight high-quality signals (e.g., cell type composition) while still leveraging informative features from the noisy modality (e.g., transcriptomic profile). Importantly, this mechanism involves nonlinear interactions between the modalities, facilitating the discovery of cross-modal features that may be uninformative in isolation. In Scenario iii, where the transcriptomic profile remains high SNR and cell type composition is degraded, we observed the opposite trend (**Figure 3G**). GraphScrDom_CT’s performance decreased substantially (ARI = 0.555), while GraphScrDom maintained strong performance with an ARI of 0.875 (**Figure 3H**), again outperforming all other models across evaluation metrics (**Figure 3I**). These results further highlight GraphScrDom’s ability to shift emphasis toward the more informative modality under varying SNR conditions.

Overall, the simulation study demonstrates that even with limited scribble annotations, GraphScrDom achieves high spatial domain detection accuracy using either a single modality or both. While either transcriptomic profiles or cell type compositions alone can provide informative cues for domain segmentation, their combination yields consistently better performance. When one modality is degraded by noise, GraphScrDom remains robust by leveraging the adaptive gated fusion to prioritize the informative modality and denoise the less reliable one. This design ensures both the robustness and effectiveness of GraphScrDom across diverse scenarios in spatial domain detection.

### GraphScrDom improved the identification of known anatomical layers of human dorsolateral prefrontal cortex 10x Visium data

We evaluated GraphScrDom on the Human DLPFC ST dataset, which includes 12 adult brain tissue sections collected from three donors and profiled using the 10x Visium platform. Specifically, samples #151507–#151510 are from donor 1, #151569–#151572 from donor 2, and #151673–#151676 from donor 3, with sections from each donor arranged sequentially along the anterior–posterior axis. Each sample contains between 3,460 and 4,789 spatial spots and measures the expression of 33,538 genes. Anatomical annotations for six cortical layers and the white matter region are provided based on the original study (Maynard et al., 2021). To estimate cell type compositions, we utilized single-nucleus RNA-seq (snRNA-seq) data from two donors in the same study, profiled using the 10x Chromium platform. The reference dataset consists of 77,581 cells with expression measured across 25,321 genes and annotated into 25 cell types. The intersection of genes between the ST and scRNA-seq datasets was used as input to Cell2location (Kleshchevnikov et al., 2022) for deconvolution, producing spot-level cell type composition estimates. These compositions, along with spatial gene expression profiles, were used as input to GraphScrDom for tissue segmentation. For all comparison methods, we followed preprocessing steps consistent with their respective original studies. Additionally, to ensure a fair comparison, GraphScrDom and ScribbleDom were both trained using the same set of scribble annotations previously generated for the ScribbleDom study except slice #151510. We reannotated slice #151510 to match the scope of scribble annotation for the other samples. These annotations cover between 3.98% and 8.03% of spots across the 12 samples. Further implementation details are provided in the Methods section.

After applying GraphScrDom and competing methods to identify spatial domains, we compared their results to the ground truth annotations. As an illustrative example, we presented the segmentation outcomes for four tissue slices from donor 3 (#151673–#151676) (**Figure 4A, Supplementary Figure S1)**. Visual inspection reveals that GraphScrDom consistently produces spatial domains that are well-aligned with the annotated cortical layers across all slices. In contrast, Leiden clustering exhibits the weakest performance, successfully identifying only the white matter (WM), while the remaining clusters are mixed among other layers. This suggests that conventional single-cell clustering approaches, which do not incorporate spatial context, may be unsuitable for spatial domain detection. Similar to Leiden, SpaGCN accurately identifies WM across all four slices but struggles to separate its boundary from Layer 6 and performs poorly in delineating other cortical layers. ScribbleDom, GraphST, and IRIS all produce overall layer-like structures, though their performance varies across slices. ScribbleDom consistently recovers Layer 1, Layer 2, and WM across all slices, with Layer 3 recovered in slices #151673 and #151674, while Layers 4 and 5 appear mixed. GraphST performs well in identifying layers for slice #151673, but its layer boundaries become increasingly ambiguous in other slices. For slice #151674, GraphST shows better separation of WM and Layers 1–3, yet Layers 4–6 are not correctly distinguished. In slices #151675 and #151676, GraphST accurately identifies WM and Layers 4–6 but misclassifies Layers 1–3. IRIS produces smooth layer structures that resemble the annotated shapes but exhibits incorrect layer thickness. Other methods, such as STAGATE and BayesSpace, show considerable variability in performance. STAGATE successfully identifies well-defined layers only for slice #151673, with poor segmentation on the remaining slices. BayesSpace produces clear separation for slices #151673 and #151675 but fails to recover consistent boundaries for the other two slices, often generating ragged or irregular layer demarcations.

**Figure 4.**
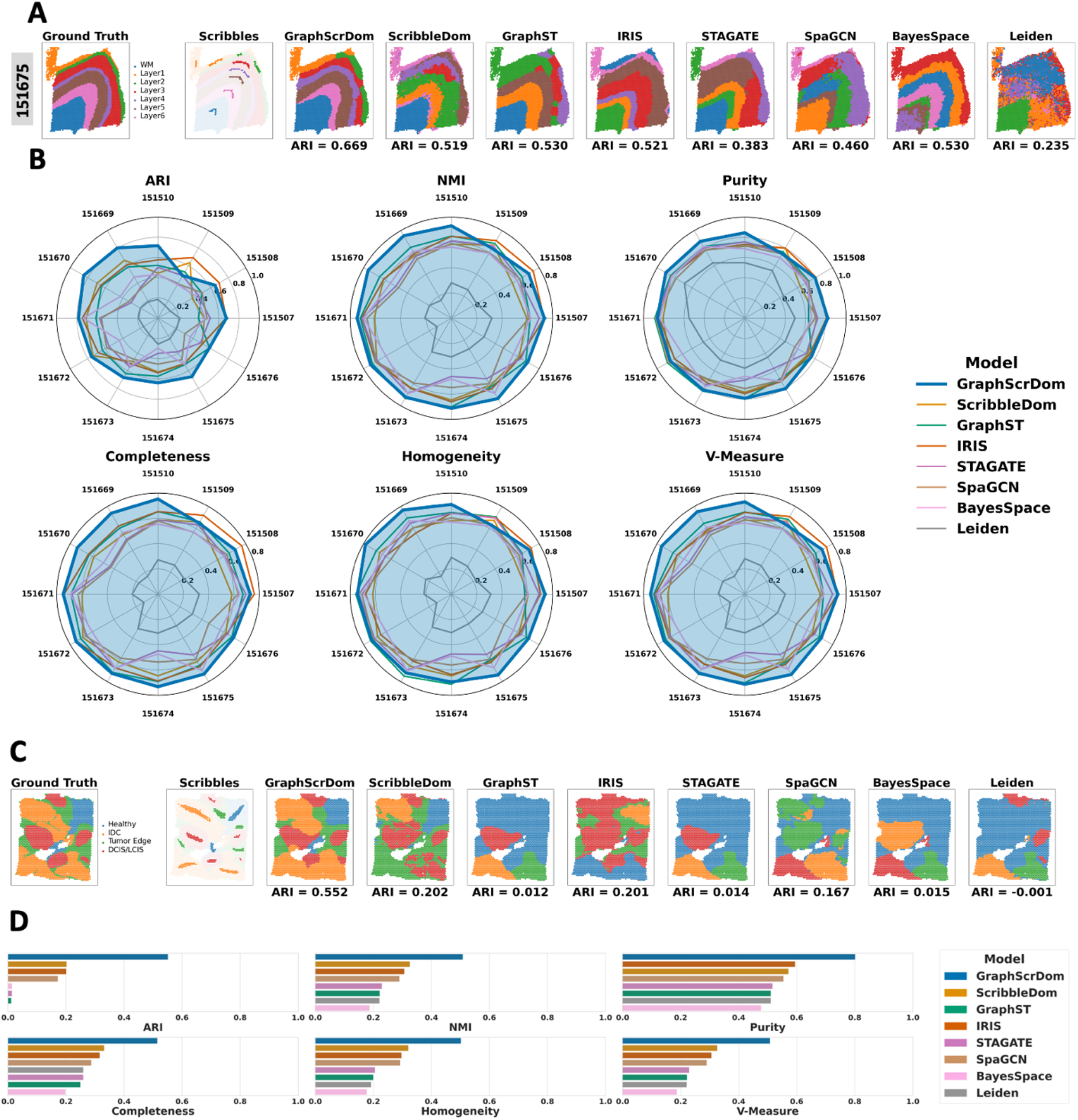
Evaluation of GraphScrDom on Visium spatial transcriptomics datasets. **A**. Spatial domain detection results on #151675 from the Human DLPFC dataset. The first two columns display ground truth annotations and scribble inputs. The remaining columns show predicted domains from GraphScrDom, ScribbleDom, GraphST, IRIS, STAGATE, SpaGCN, BayesSpace, and Leiden. **B**. Quantitative comparison of model performance across all 12 DLPFC samples using six evaluation metrics: Adjusted Rand Index (ARI), Normalized Mutual Information (NMI), Purity, Completeness, Homogeneity, and V-Measure. **C**. Spatial domain detection on the Human Breast Cancer dataset. Ground truth annotations and scribble inputs are shown alongside predictions from each method. **D**. Evaluation metrics on the breast cancer dataset.

We further benchmarked all methods on four additional slices (#151669–#151672) from donor 2 (**Supplementary Figure S1)**. Unlike the samples from donor 1, these slices lack Layer 1 and Layer 2 but exhibit an expanded region corresponding to Layer 3. Despite this variation, GraphScrDom successfully recovered smooth and clearly delineated layer structures across all slices. In contrast, Leiden again demonstrated the weakest performance, failing to separate any distinct layers. Other competing methods—including ScribbleDom, GraphST, IRIS, STAGATE, and BayesSpace—were able to detect layer-like structures but showed notable limitations. ScribbleDom produced layer boundaries that closely followed the ground truth, although the boundaries appeared ragged. IRIS identified distinct layers; however, most were misaligned with the annotated ground truth. GraphST, STAGATE, and BayesSpace struggled in separating the large Layer 3 region, which was frequently mixed with Layer 4, leading to reduced segmentation fidelity.

A unique “sandwich” layer topology was observed in slices #151507–#151510 from donor 3 (**Supplementary Figure S1**), presenting a significant challenge for spatial domain detection methods. In these samples, identical layers appear in non-contiguous locations, separated by other layers. For example, two thin regions of Layer 2 are split by a wide intervening Layer 1, and in a more complex case, two thick Layer 3 regions are separated by two thin Layer 2 regions with a broad Layer 1 segment in between. This complex structure requires precise modeling of spatial continuity and domain boundaries. GraphScrDom and IRIS both successfully segmented all layers, including the sandwich-like structures, with predicted layer thicknesses closely matching the manual annotations. ScribbleDom also recovered the overall structure, but the boundaries between layers appeared more blurred. In contrast, Leiden performed poorly, merging all layers into a single mixed domain, while BayesSpace failed to recover most layers altogether. SpaGCN did not recover Layer 2 in any of the slices and generally failed to produce well-demarcated boundaries. GraphST performed well in separating the sandwich-like layers in slices #151509 and #151510, successfully identifying both Layer 2 and Layer 3. However, it tended to overestimate the thickness of Layer 2 and underestimate Layer 3 compared to the ground truth. STAGATE showed a similar pattern, recovering Layers 2 and 3 in slices #151507 and #151509, but with poor separation of other layers. This comprehensive benchmark across Human DLPFC slices—capturing variability in layer topology across three donors and multiple samples—demonstrates the robustness of GraphScrDom in accurately detecting spatial domains under complex anatomical configurations.

To quantitatively assess performance, we used the widely adopted Adjusted Rand Index (ARI) as the performance evaluation metric. GraphScrDom achieved the highest median ARI of 0.674 and ranked first in 11 out of 12 samples from the Human DLPFC data (**Figure 4B**). On average, GraphScrDom improved ARI by 13% to 268% compared to competing methods (**Supplementary Table S9**). The improvement was also statistically significant based on the Wilcoxon signed-rank test (p-value < 0.05). In addition to ARI, we evaluated five other metrics— Normalized Mutual Information (NMI), Purity, Completeness, Homogeneity, and V-Measure. GraphScrDom consistently performed well across all metrics, further supporting its robustness and generalizability in spatial domain detection (**Figure 4B and Supplementary Table S3-S8**).

### GraphScrDom accurately identifies spatial domains on Human Breast Cancer Tissue 10x Visium data

We further evaluated GraphScrDom on a 10x Visium dataset from a human breast cancer tissue slice, which profiles the expression of 36,601 genes across 3,798 spatial spots. This sample includes four annotated tissue regions—ductal/lobular carcinoma in situ (DCIS/LCIS), invasive ductal carcinoma (IDC), healthy tissue, and tumor edge—based on expert pathological annotation. To estimate spot-level cell type composition, we used the processed single-cell RNA-seq data from GSE176078 (Wu et al., 2021), which contains 100,064 cells and 29,067 genes across 18 annotated cell types. For GraphScrDom and ScribbleDom, scribble annotations covering approximately 10% of the spots were manually generated using the GraphScrDom GUI toolkit.

Unlike the Human DLPFC dataset, which exhibits a clear layer-like spatial organization, the breast cancer tissue slice lacks such structured layering. Spatial domains are often noncontiguous and dispersed across different regions of the tissue. For instance, DCIS/LCIS appears in four disjoint areas—top, bottom, left, and right—posing significant challenges for both computational domain detection and manual contour-based segmentation by pathologists. Under these complex spatial conditions, GraphScrDom accurately identified all four spatial domains in alignment with expert annotations, leveraging the sparse supervision provided by scribble labels. In contrast, other methods struggled to recover the correct spatial structures. ScribbleDom produced noisier predictions with less regional continuity and, notably, misclassified a large IDC region in the bottom of the slice as DCIS/LCIS. GraphST, STAGATE, BayesSpace, and Leiden exhibited similar failure patterns, incorrectly merging multiple regions from the healthy tissue, IDC, and tumor edge into a single large cluster in the upper and central parts of the tissue. While SpaGCN and IRIS identified more fragmented regions, their predicted domains did not align well with the pathologist’s annotations. These observations are further supported by quantitative metrics. GraphScrDom achieved the highest ARI of 0.552 among all competing methods (**Supplementary Table S3**). It also outperformed others across five additional evaluation metrics—NMI, Purity, Completeness, Homogeneity, and V-Measure—as shown in **Figure 4D** and **Supplementary Tables S3–S8**.

### GraphScrDom provides robust spatial domain detection on Mouse Frontal Cortex MERFISH data

In addition to benchmarking on the 10x Visium platform, we evaluated GraphScrDom on spatial transcriptomics data generated at single-cell resolution. Specifically, we used the Mouse Frontal Cortex dataset profiled using the MERFISH platform, which includes 9 samples, each derived from a different donor. Each sample contains between 21,002 and 40,484 cells and was measured using a consistent panel of 374 genes. All samples are annotated with 8 anatomical brain regions, including corpus callosum, pia mater, striatum, olfactory region, brain ventricle, and cortical layers II/III, V, and VI (Allen et al., 2023). To support weak supervision, we performed scribble annotations on each sample, covering between 3.81% and 9.52% of cells per slice. To derive spot-level cell type composition, we paired each MERFISH sample with an age-matched reference snRNA-seq dataset from the same study. Samples from donors 1, 4, 7, and 8 (4-week-old mice) were matched to a reference snRNA-seq dataset profiling 20,984 genes across 21,728 cells, annotated with 13 cell types. Samples from donors 2, 3, 5, 6, and 9 (90-week-old mice) were matched to a corresponding 90-week-old snRNA-seq dataset, also profiling 20,984 genes across 20,152 cells with the same 13 annotated cell types.

We compared the spatial domain detection results across benchmarked methods (**Figure 5A, Supplementary Figure S2)**. BayesSpace and ScribbleDom were excluded from this comparison, as both are specifically designed for grid-based ST platforms (e.g., 10x Visium) and are not directly applicable to single-cell resolution datasets such as MERFISH. When the layer-like tissue structure is well-defined, as observed in samples from donors 1, 3, 7, 8, and 9, GraphScrDom consistently outperforms other methods, accurately segmenting spatial domains with smooth and anatomically aligned boundaries. For donor 3, GraphScrDom clearly separates all annotated brain regions with high boundary fidelity. While GraphST, IRIS, and STAGATE also achieve reasonable segmentation of large regions such as the corpus callosum and striatum, their ability to distinguish between cortical layers V, VI, and II/III is limited by fuzzy or overlapping boundaries. SpaGCN and Leiden perform worse, showing significant mixing of the three cortical layers. Notably, IRIS uniquely detects the pia mater, which is missed by other methods. In donor 7, all methods are able to identify major regions such as the corpus callosum, striatum, and pia mater. However, only GraphScrDom, GraphST, and STAGATE accurately segment all eight brain regions with smooth and well-defined boundaries. IRIS partially captures the cortical layers but with jagged and inconsistent borders, while SpaGCN and Leiden again fail to clearly separate cortical layers V, VI, and II/III. For donor 8, GraphScrDom successfully detects all annotated regions with clear and continuous boundaries. STAGATE is able to identify the three cortical layers but performs poorly in segmenting the striatum, which is erroneously merged with adjacent regions. Other methods either misclassify the cortical layers entirely or yield poorly defined region boundaries. In samples where the spatial layer structure is less apparent and domain boundaries become more diffuse (donors 2, 4, 5, and 6), performance declines across all methods. Nevertheless, GraphScrDom—along with GraphST—continues to lead in performance, highlighting its robustness even in challenging spatial contexts lacking clear anatomical demarcations.

**Figure 5.**
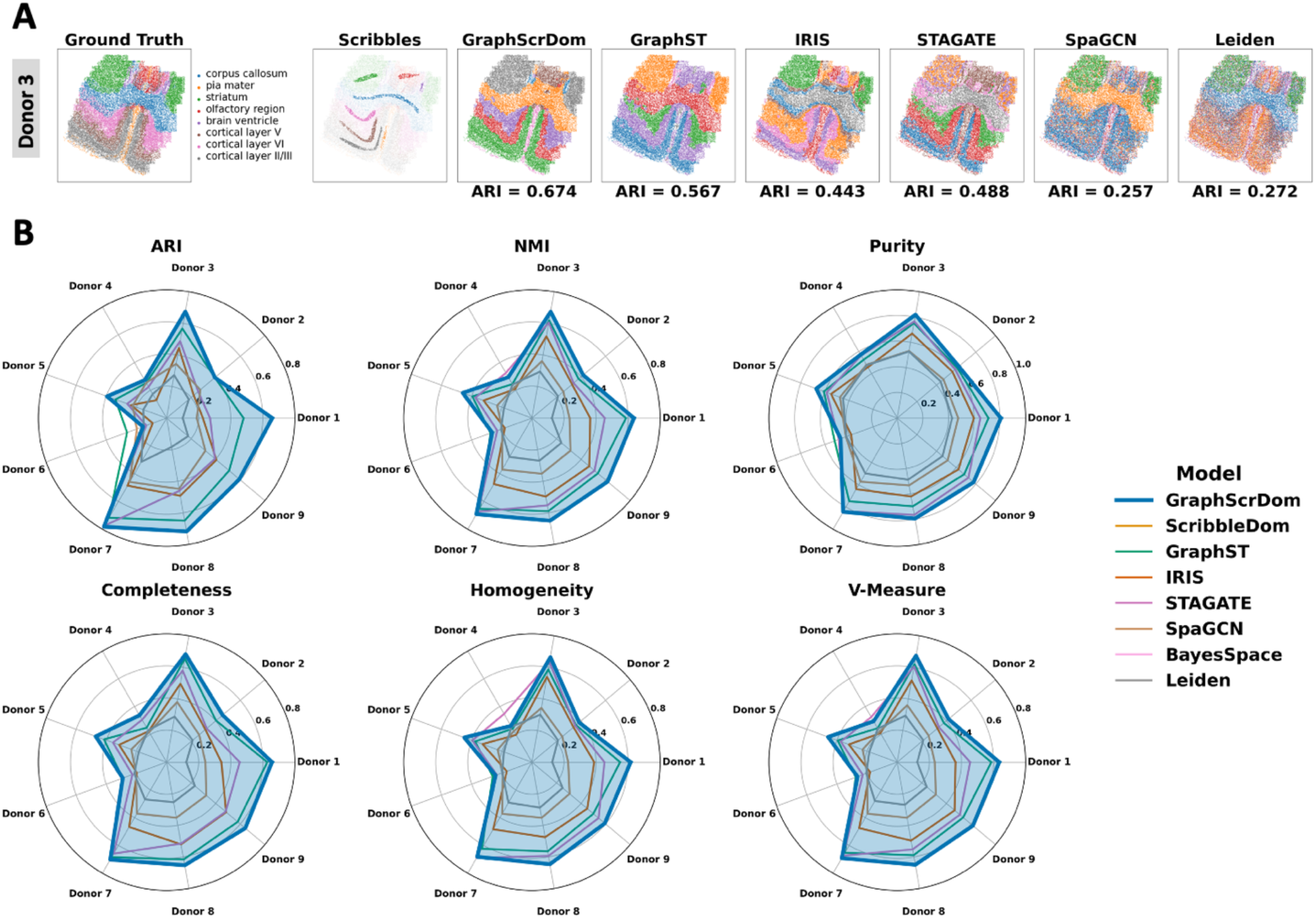
Evaluation of GraphScrDom on the MERFISH Mouse Frontal Cortex dataset. **A**. Spatial domain detection results for donors 3. The first two columns display ground truth annotations and scribble inputs. The remaining columns show predicted spatial domains generated by GraphScrDom and other benchmark methods. **B**. Quantitative evaluation across all nine MERFISH donors using six performance metrics: Adjusted Rand Index (ARI), Normalized Mutual Information (NMI), Purity, Completeness, Homogeneity, and V-Measure.

In addition to spatial visualizations, we quantitatively evaluated model performance using ARI and five additional metrics across all nine MERFISH samples. GraphScrDom achieved the highest median ARI of 0.593, ranking first in 8 out of 9 samples (**Figure 5B** and **Supplementary Table S3**). The second-best performer, GraphST, attained a median ARI of 0.480. Compared to other benchmark methods, GraphScrDom improved ARI scores by 8.5% to 190%. Statistical significance was confirmed using Wilcoxon signed-rank tests, with p-values < 0.05 across all pairwise comparisons (**Supplementary Table S10**). Beyond ARI, GraphScrDom also outperformed competing methods across the remaining evaluation metrics—NMI, Purity, Completeness, Homogeneity, and V-Measure—as shown in **Figure 5B** and **Supplementary Tables S3–S8**. These results further demonstrate the versatility and robustness of GraphScrDom, which generalizes effectively across both bulk and single-cell resolution data, and is adaptable to both grid-based and non-grid-based spatial transcriptomics platforms.

### GraphScrDom achieves accurate spatial domain detections across diverse single cell spatial platforms

To further assess the generalizability of GraphScrDom, we extended our evaluation to three single-cell resolution spatial transcriptomics (ST) datasets derived from different mouse brain regions and profiled using three distinct platforms: osmFISH, BaristaSeq, and STARmap. These datasets encompass a range of spatial resolutions, gene coverages, and anatomical complexities, providing a comprehensive testbed for validating the adaptability of GraphScrDom. The Mouse Somatosensory Cortex dataset, generated using the osmFISH platform, includes 4,839 spatial locations and measures expression for 33 genes across 11 annotated brain regions, including Layer 2/3 lateral, Layer 2/3 medial, Layer 3/4, Layer 4–6, hippocampus, ICC, pia/layer 1, ventricle, and white matter (Codeluppi et al., 2018). The Mouse Primary Visual Cortex dataset, profiled using BaristaSeq, contains 4,432 spots and 76 genes across six annotated spatial domains, including L1, L2/3, L4–6, and LWM (Chen et al., 2015). The Mouse Visual Cortex dataset, generated using the STARmap platform, comprises 1,207 spots and 1,020 genes across seven annotated domains, including L1, L2/3, L4–6, ICC, and HPC (Wang et al., 2018). To derive cell type compositions, we used age- and region-matched scRNA-seq reference datasets. For the osmFISH and BaristaSeq datasets, we adopted scRNA-seq data from a study focused on regional profiling of the mouse isocortex, including primary visual cortex and primary somatosensory cortex, generated using the SMART-seq2 protocol (Yao et al., 2021).

Region-specific subsets were extracted based on anatomical annotations corresponding to the target ST tissues. The reference for the Mouse Somatosensory Cortex consists of 7,531 cells and 35,433 genes across 21 cell types, while the reference for the Mouse Primary Visual Area includes 15,572 cells and 35,433 genes across 23 cell types. For the STARmap dataset, we utilized an scRNA-seq reference from a study investigating gene expression changes in the mouse visual cortex in response to sensory experience (Hrvatin et al., 2018; GEO accession: GSE185862). This dataset includes 2,219 cells and 25,202 genes, clustered into 31 annotated cell types. Scribble annotations were manually generated for each ST dataset to support weak supervision, covering between 10% and 12% of spatial locations per dataset (**Supplementary Table S1**). A comprehensive summary of all ST and reference scRNA-seq datasets is provided in **Supplementary Tables S1** and **S2**.

We excluded BayesSpace and ScribbleDom from this analysis, as both methods are designed specifically for grid-based platforms and are not applicable to single-cell resolution ST datasets. Notably, the Mouse Somatosensory Cortex ST dataset (osmFISH) profiles only 33 genes per spot, which limits the ability to capture transcriptional heterogeneity and makes it challenging to distinguish spatial domains with distinct expression patterns. Despite this limitation, GraphScrDom successfully delineates all annotated spatial domains with smooth and coherent region boundaries that closely align with the manual annotations (**Figure 6A**). Graph-based approaches such as GraphST, STAGATE, SpaGCN, and Leiden partially recover the layered structure, reliably identifying regions such as the hippocampus, ICC, white matter, Layer 6, and Pia Layer 1, but fail to separate Layers 2–5. IRIS performs the worst, failing to detect most layers. Quantitatively, GraphScrDom achieves the highest ARI of 0.723, and consistently outperforms all competing methods across all evaluation metrics (**Figure 6B** and **Supplementary Table S4**).

**Figure 6.**
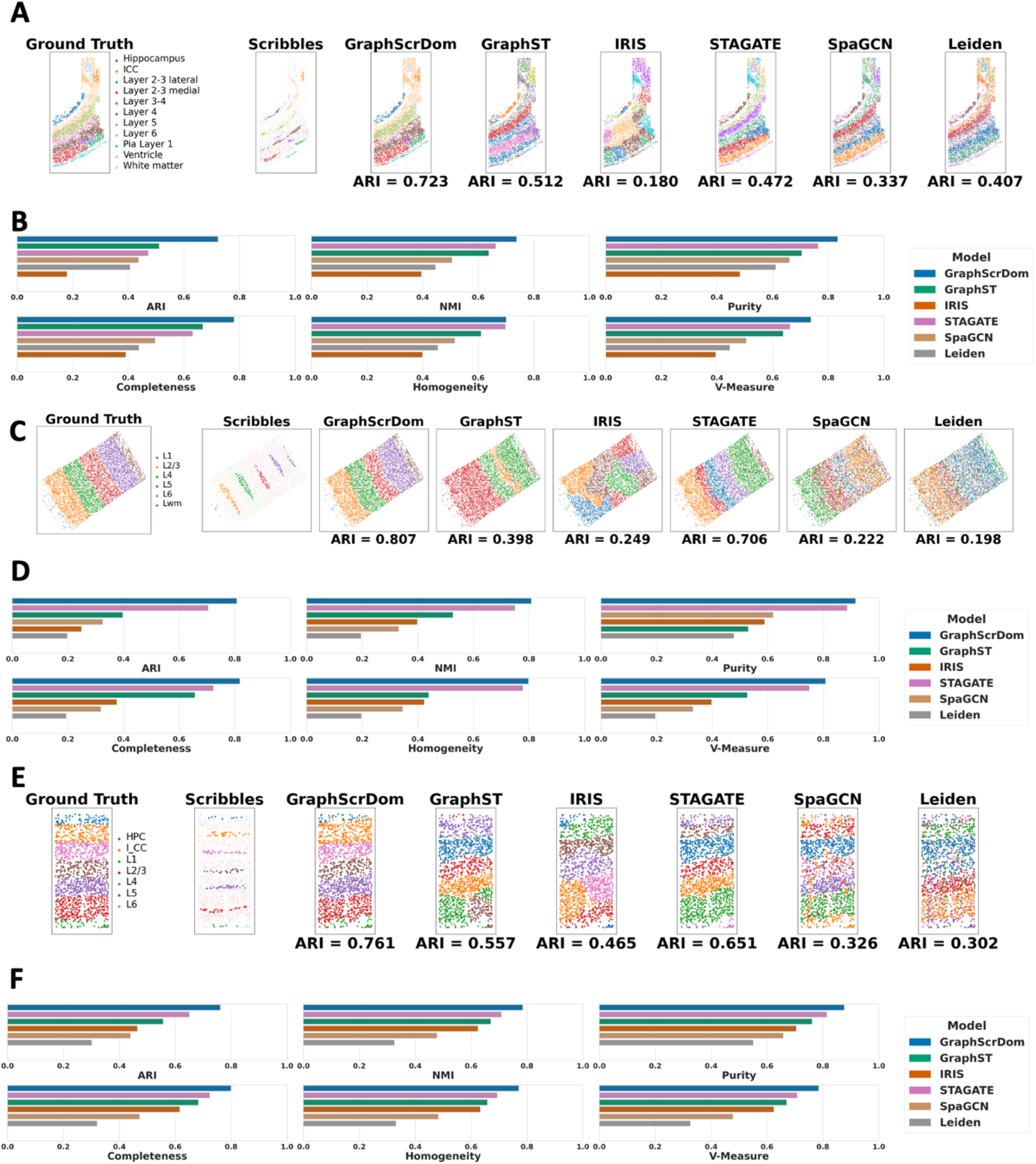
Evaluation of GraphScrDom on single-cell spatial transcriptomics (ST) datasets. **A**. Spatial domain detection results on the osmFISH Mouse Somatosensory Cortex dataset. The first two columns display ground truth annotations and scribble inputs. The remaining columns show predicted spatial domains generated by GraphScrDom and other benchmark methods. **B**. Quantitative evaluation on the osmFISH dataset. **C**. Spatial domain detection results on the BaristaSeq Mouse Primary Visual Cortex dataset. **D**. Evaluation metrics on the BaristaSeq dataset. **E**. Spatial domain detection results on the STARmap Mouse Visual Cortex dataset. **F**. Evaluation metrics on the STARmap dataset.

The Mouse Primary Visual Area ST dataset (BaristaSeq) also profiles a small gene panel (76 genes), but includes six well-defined cortical layers with clear boundaries. In this relatively easier setting, only GraphScrDom and STAGATE successfully separate all layers (**Figure 6C**), whereas other methods fail to recover the domain structure accurately. GraphScrDom achieves the highest ARI of 0.807, followed by STAGATE (ARI = 0.706). This performance trend holds across other metrics as well (**Figure 6D** and **Supplementary Tables S3–S8**).

The Mouse Visual Cortex ST dataset (STARmap), like BaristaSeq, also targets the visual cortex, but profiles a substantially larger gene panel (1,020 genes). This increased gene coverage enhances the ability to resolve spatial domains, leading to improved performance for most methods. For example, ARI scores notably increase for IRIS (0.465 vs. 0.249), GraphST (0.557 vs. 0.398), SpaGCN (0.326 vs. 0.222), and Leiden (0.302 vs. 0.198) compared to BaristaSeq, highlighting the benefit of transcriptomic richness for spatial domain detection. Both GraphScrDom and STAGATE show stable performance across the two platforms, demonstrating robustness to differences in gene coverage. GraphScrDom again outperforms all methods with the highest ARI of 0.761, followed by STAGATE (ARI = 0.651) (**Figure 6E**). Across all additional metrics, GraphScrDom maintains a substantial lead (**Figure 6F** and **Supplementary Tables S3– S8**). Taken together, these results from three distinct single-cell resolution ST datasets highlight the robustness and generalizability of GraphScrDom. It consistently achieves top performance across a variety of experimental conditions—regardless of platform, gene panel size, or spatial domain complexity.

## Conclusion and Discussion

Spatial domain detection is a critical task in spatial transcriptomics, enabling the identification of anatomical and functional tissue regions. Most existing approaches rely solely on transcriptomic heterogeneity—typically driven by highly variable genes (HVGs) or spatially variable genes (SVGs)—to characterize tissue structure (Sun et al., 2020; Svensson et al., 2018; Weber et al., 2023). Recent studies have shown that cell type composition exhibits distinct enrichment patterns across anatomical regions (Biancalani et al., 2021; Kleshchevnikov et al., 2022), highlighting the potential of incorporating cell type composition to enhance spatial domain detection. This hypothesis has been further supported by IRIS (Ma & Zhou, 2024), which integrates both cell type composition and transcriptomic profiles to improve tissue segmentation. In parallel, computer vision techniques for image segmentation have adopted weakly supervised learning strategies, such as using sparse scribble annotations (e.g., freeform strokes), followed by semi-supervised learning to achieve full-scale segmentation. These strategies alleviate the burden of detailed polygon annotations, which are time-consuming and prone to error due to morphological variability and irregular tissue structures.

Here, we present GraphScrDom, a hybrid multimodal graph neural network (GNN) model that integrates spatial transcriptomic data and cell type composition for spatial domain detection. GraphScrDom employs gene expression and cell type composition encoder-decoders within an end-to-end framework, jointly learning spot embeddings and predicting spatial domain labels. Unlike ScribbleDom, which uses convolutional neural networks (CNNs), GraphScrDom utilizes a GNN-based classifier, making it applicable to both grid-based (e.g., 10x Visium) and non-grid-based (e.g., MERFISH) ST platforms. Furthermore, the multimodal framework leverages a gated mechanism to adaptively balance the contributions of transcriptomic and cell type information, improving robustness. A user-friendly software toolkit is provided, including an annotation module for scribble input and a training module for semi-supervised domain prediction.

We extensively benchmarked GraphScrDom using both simulations and real-world ST datasets. In simulations, GraphScrDom was evaluated under three scenarios: (i) both modalities informative, (ii) presence of high noise in gene expression and (iii) presence of noise in cell type composition. In all cases, GraphScrDom exhibited robust performance and showed a clear advantage in integrating complementary signals. For real data applications, GraphScrDom was compared against seven state-of-the-art methods—ScribbleDom, GraphST, IRIS, STAGATE, SpaGCN, BayesSpace, and Leiden—across six evaluation metrics (ARI, NMI, Purity, Completeness, Homogeneity, and V-Measure) on six spatial transcriptomics datasets from five platforms, varying in spatial resolution (bulk or single-cell), gene coverage, and sample size. GraphScrDom consistently achieved strong performance across these datasets. On the Human DLPFC ST dataset (10x Visium), which includes 12 slices from 3 donors, GraphScrDom accurately captured annotated cortical layers, including complex “sandwich” configurations. On the single-cell resolution Mouse Frontal Cortex dataset (MERFISH), GraphScrDom outperformed all competitors in delineating fine-grained structures across 9 donors. These results demonstrate GraphScrDom’s robustness across spatial resolutions, tissue architectures, and donor variability. In a challenging example, the Human Breast Cancer ST dataset (10x Visium) contains highly heterogeneous tissue where a domain may span multiple disconnected regions. GraphScrDom effectively recovered these domains, underscoring the value of weak supervision via scribble annotations, particularly when unsupervised methods fail to capture complex structure. We further evaluated GraphScrDom on ST datasets from three additional single-cell resolution platforms—osmFISH (Mouse Somatosensory Cortex), BaristaSeq (Mouse Primary Visual Cortex), and STARmap (Mouse Visual Cortex)—each featuring low to moderate gene coverage. Despite limited transcriptomic input, GraphScrDom continued to outperform alternatives by leveraging scribble supervision and scRNA-seq-derived cell type compositions.

While GraphScrDom demonstrates strong performance, it currently relies on expert-provided scribble annotations, which may not always be readily available in practice. Additionally, the number of spatial domain classes must be predefined, which constrains the model’s capacity to discover novel or unexpected domains. As such, GraphScrDom is best suited for targeted domain detection rather than fully exploratory analyses. Several extensions are planned to enhance GraphScrDom’s flexibility and applicability. In the absence of scribble annotations, we propose an unsupervised initialization strategy in which a joint embedding is learned in an end-to-end manner, followed by unsupervised clustering. The resulting cluster centers can then serve as pseudo-scribble annotations to facilitate weakly supervised domain detection. Beyond transcriptomic profiles and cell type composition, future versions of GraphScrDom will also incorporate tissue morphology captured from histology images. This can be achieved by integrating visual features extracted from deep learning models pre-trained on large histopathology datasets (Huang et al., 2023; Lu et al., 2024), further enriching the multimodal representation. Moreover, the current implementation is designed for single-slice analysis. To enable multi-slice spatial domain detection, we plan to integrate image registration algorithms to reconstruct a 3D tissue architecture across serial sections. In this setting, a unified graph structure can be constructed, and scribble annotations from one or more slices can guide weakly supervised learning to infer spatial domains across all slices.

## Methods

### Data preprocessing

We benchmarked GraphScrDom and other competing methods on six published spatial transcriptomics (ST) datasets for the task of spatial domain detection. These datasets encompass diverse tissue types (e.g., visual cortex, frontal cortex, breast), multiple species (e.g., human and mouse), and various ST platforms, including both bulk-resolution (e.g., 10x Visium) and single-cell resolution technologies (e.g., STARmap, osmFISH, BaristaSeq) (**Supplementary Table S1**). For each ST dataset, a corresponding reference single-cell RNA-seq (scRNA-seq) dataset from the same tissue, with cell type annotations, is used for downstream integration (**Supplementary Table S2**). As GraphScrDom requires spatial gene expression, spatial coordinates, and cell type compositions as input, we applied standardized data preprocessing procedures to normalize expression data and derive cell type compositions. Specifically, spatial gene expression values were first scaled by total count per spot, multiplied by a scaling factor of 10,000, and then log-transformed. To minimize the effect of extreme values, expression values were clipped at a maximum of 10. From this processed matrix, the top 3,000 highly variable genes (HVGs) were selected. For ST datasets containing fewer than 3,000 genes, all available genes were retained. To infer cell type composition, we adopted Cell2location (Kleshchevnikov et al., 2022), a Python-based deconvolution method that has demonstrated strong performance across multiple benchmarking studies (Gaspard-Boulinc et al., 2025; Li et al., 2023). Its seamless integration into our Python-based framework further facilitated its use. Cell2location requires as input: (i) the ST gene expression matrix for the set of genes shared between ST and scRNA-seq data, and (ii) the scRNA-seq reference data, including cell type annotations and the single-cell gene expression matrix. Following Cell2location’s recommended preprocessing, we apply minimal gene filtering to the scRNA-seq data by retaining genes that are expressed in more than five cells, present in over 5% of cells, and have a non-zero mean expression greater than 1.12. The output from Cell2location includes the estimated abundance of each cell type at every spatial location. For spot-level ST platforms (e.g., 10x Visium), the estimated cell counts per type within each spot are reported, which we normalize into cell type proportions. For cell-level ST platforms (e.g., STARmap), Cell2location estimates cell type probabilities per cell. The resulting cell type composition, together with the spatial gene expression matrix from the top 3,000 HVGs, is used as model input for GraphScrDom. For other methods, we followed their original preprocessing protocols or recommend procedures.

### GraphScrDom model

#### Graph Construction

As GraphScrDom is built upon a graph neural network, we begin by constructing the spatial graph based on the coordinates of each spot or cell, applicable to both spot and single-cell resolution platforms. Specifically, we define an undirected graph *G* = (*V, E*), where *V* denotes the set of spatial locations (spots or cells) and *E* represents the edges connecting neighboring locations. To define the edges, each node *v*_*i*_ ∈ *V* is connected to its *K* nearest neighbors based on Euclidean distance in the spatial coordinate space. This results in a symmetric adjacency matrix *A* ∈ ℝ^*N*×*N*^, where a_*i,j*_ = a_*j,i*_ = 1 if nodes *v*_*i*_ and *v*_*j*_ are connected, and a_*i,j*_ = a_*j,i*_ = 0 otherwise. The number of neighbors, *K*, is a tunable hyperparameter. In our experiments, GraphScrDom achieves optimal performance when *K* is set to 20.

#### Overview of GraphScrDom

GraphScrDom is a hybrid multimodal graph neural network that incorporates three reconstruction modules and a classification module. These modules are designed to jointly leverage spatial gene expression and cell type composition for spatial domain detection. The three reconstruction modules are described as follows: (i) Gene expression encoder-decoder: The original gene expression profile *x*_*gi*_ for spot *s*_*i*_ is encoded to the latent embedding as *z*_*gi*_, which is then used to reconstruct the gene expression profile of the same spot, yielding 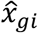. (ii) Cell type composition encoder-decoder: The original cell type composition profile *x*_*ci*_ for spot *s*_*i*_ is encoded to the latent embedding as *z*_*ci*_, which is then used to reconstruct the gene expression of the same spot, resulting in 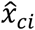. (iii) Hybrid encoder-decoder: The latent embeddings *z*_*gi*_ and *z*_*ci*_ are concatenated and passed through a dynamic gated fusion mechanism to generate a hybrid embedding *z*_*fi*_. This fused embedding is used to reconstruct the gene expression profile 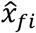, and also serves as the input of the classifier for spatial domain prediction.

#### GCN-based encoder-decoder for learning spot latent representation

We employ two separate Graph convolution network (GCN)-based encoders (Kipf & Welling, 2017) to learn spot-level latent representations: one for the gene expression profile and one for cell type composition, both incorporating spatial neighborhood information.

Each encoder takes as input the spatial adjacency matrix and the respective feature matrix (gene expression or cell type composition), and outputs a latent representation, which is then passed through a symmetric GCN decoder to reconstruct the original input. We use the gene expression encoder-decoder as an illustrative example. Let the spatial gene expression matrix be denoted by *X*_*g*_ ∈ ℝ ^*N*×*m*^, where *N* is the number of spots and *m* is number of highly variable genes. Let *A* ∈ ℝ ^*N*×*N*^ denote the symmetric adjacency matrix constructed from spatial coordinates, and define Ã = *A* + *I*_*N*_, where *I*_*N*_ is the identity matrix. The corresponding degree matrix is 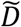 ^with entries 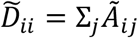^_. The latent representation *Z*_*g*_ ∈ ℝ ^*N*×*d*^ is obtained via a GCN_ encoder with *L* layers. The (*l*+1)-th GCN layer is defined as:

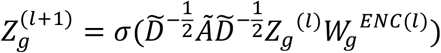

where σ(·) is the ReLU activation function, *W*_*g*_^*ENC*(*l*)^is a layer-specific trainable weight matrix in the encoder, *Z*_*g*_^(0)^ = *X*_*g*_. The decoder reconstructs the input from the learned latent representation. Specifically, the decoder takes *Z*_*g*_ as the initial input, i.e., *H*_*g*_^(0)^ = *Z*_*g*_, and applies a symmetric GCN architecture to reconstruct the gene expression matrix 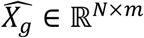. The decoder layer at depth (*l*+1)-th is defined as:

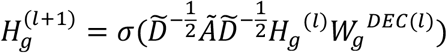

where *W*_*g*_^*DEC*(*l*)^is a layer-specific trainable weight matrix in the decoder. The final output 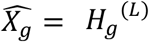 is compared with the original input to compute the reconstruction loss as:

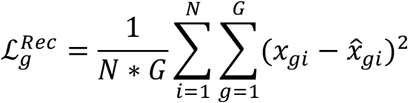

The same architecture is applied to the cell type composition encoder-decoder. Let *X*_*c*_ ∈ ℝ ^*N*×*C*^ denote the cell type composition matrix (with *C* cell types). The latent representation *Z*_*c*_ is used as the first decoder input, i.e., *H*_*c*_^(0)^ = *Z*_*c*_, and the final output 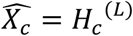 is compared with the original input to compute the reconstruction loss as:

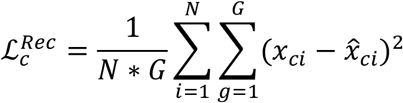

In our implementation, we use a single GCN layer (*L* = 1) for both the encoder and decoder, which we find sufficient to capture local structure and achieve strong performance across datasets.

#### Self-supervised contrastive learning for representation refinement

To enhance the discriminability of latent representations, we adopt a self-supervised contrastive learning strategy (SCL) (Veličković, 2019) into each GCN-based encoder, resulting in a SCL-GCN encoder. This approach aims to refine embeddings by encouraging the model to distinguish between meaningful (positive) and perturbed (negative) representations.

Given the input features *X* ∈ ℝ ^*N*×*m*^ (e.g., gene expression or cell type composition), we generate a corrupted version *X*^∗^ ∈ ℝ ^*N*×*m*^ by randomly shufling the rows of *X* along the spot dimension. We then input both *X* and *X*^∗^ into the same SCL-GCN encoder to obtain latent representations:

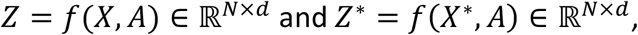

where *A* ∈ ℝ ^*N*×*N*^ is the adjacency matrix of the spatial graph. For each node *v*_*i*_, we compute it local embedding *e*_*i*_ by averaging the embedding of its *K* nearest neighbors and itself:

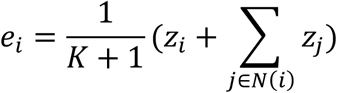

where *z*_*i*_ is the latent representation of node *v*_*i*_, and *N*(*i*) denotes the neighborhood of *v*_*i*_. The corrupted context embedding 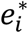 is computed analogously from *Z*^∗^. We define (*z*_*i*_, *e*_*i*_) a positive pair and 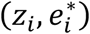 a negative pair. To discriminate between positive and negative pairs, we introduce a bilinear scoring function:

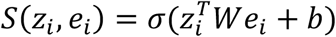

where *W* is the trainable weight matrix, *b* is the bias and σ(·) is the sigmoid function. The original contrastive loss encourages high similarity scores for positive pairs and low scores for negative pairs:

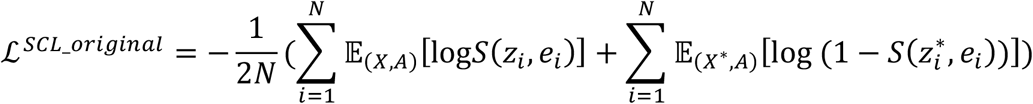

A symmetric contrastive loss is similarly defined on the corrupted embeddings:

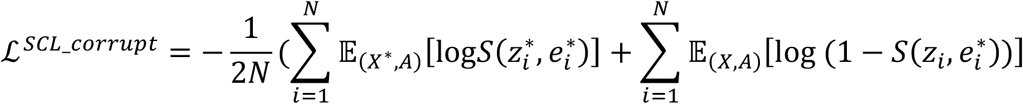

The dual contrastive loss is the sum of both the two components as:

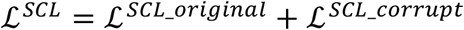

This contrastive learning loss is integrated into the overall training process. The refined representations from each encoder (gene expression and cell type composition) are subsequently passed to their corresponding decoders for reconstruction, contributing to both discriminative embedding learning and faithful input reconstruction.

#### Modality fusion with adaptive gated mechanism

In spatial transcriptomics (ST) data, gene expression captures the transcriptional landscape of each spatial spot, while cell type composition reflects the underlying cellular makeup. Integrating these two modalities can enhance representation learning by leveraging their complementary biological information. However, direct fusion may be suboptimal due to challenges such as modality-specific noise—e.g., dropout and doublets in single-cell (SC) data, or RNA diffusion and tissue artifacts in ST data (Benjamin et al., 2024)—as well as domain shifts introduced by batch effects or platform-specific differences in sample preparation and sequencing protocols (Hao et al., 2024; Cable et al., 2022). To address these challenges, we employ an adaptive gated fusion mechanism that enables data-driven weighting of the two modalities. Specifically, let *Z*_*g*_ ∈ ℝ ^*N*×*d*^ and *Z*_*c*_ ∈ ℝ ^*N*×*d*^ denote the latent representations of gene expression and cell type composition from their respective SCL-GCN encoders.

These representations are concatenated and passed through a multilayer perceptron (MLP) to compute a gating vector via

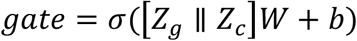

where *W* ∈ ℝ ^2*d*×*d*^, *b* ∈ ℝ ^*N*^, and σ(·) is the sigmoid function. The final fused representation is calculated as:

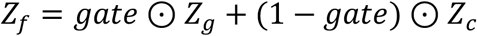

where ⊙ is element-wise multiplication. This fused embedding is refined via contrastive learning and subsequently used to reconstruct gene expression and serve as input to the classifier for spatial domain detection. The associated reconstruction loss is defined as

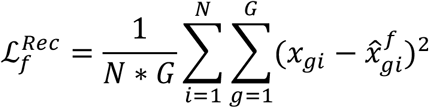

where *x*_*gi*_ is the observed gene expression and 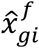 is the reconstructed value from the fused representation.

#### Semi-supervised classifier

Scribble annotations typically label only a small subset of spots, leaving the majority of spatial spots unlabeled. The objective of GraphScrDom is to infer the spatial domain labels of these unlabeled spots by leveraging the limited labeled annotations, thereby achieving comprehensive spatial domain detection. To accomplish this, GraphScrDom employs a semi-supervised graph convolutional network (GCN)-based classifier. In this framework, each node represents a spatial spot with its fused feature embedding *Z*_*f*_, and GCN leverages the spatial adjacency to propagate label information, based on the assumption that neighboring nodes in the graph are likely to share the same class. Unlike standard GCN classifiers, GraphScrDom introduces two key innovations. First, instead of decoupling representation learning and classification into separate steps, GraphScrDom trains both jointly in an end-to-end fashion, allowing embeddings to be optimized for the classification task. Second, GraphScrDom integrates both supervised and unsupervised losses into a unified objective to better utilize the unlabeled data and reduce overfitting. Specifically, the combined classification loss is defined as:

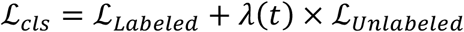

The supervised loss is a cross-entropy loss 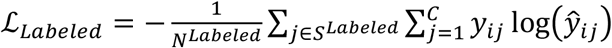 computed between prediction probability (ŷ_*ij*_) and scribble-derived ground truth labels (*y*_*ij*_) for labelled spots in each batch. *S*^*Labeled*^ is the set of labelled spots and *N*^*Labeled*^is the number of labelled spots in each batch. *N*^*Labeled*^=|*S*^*Labeled*^| and *C* is the number of classes. The unsupervised loss is another cross-entropy loss 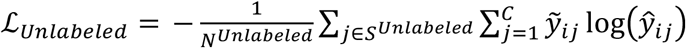 computed between prediction probability (ŷ_*ij*_) and pseudo labels 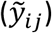 for unlabeled spots in each batch. *S*^*Unlabeled*^ is the set of labelled spots and *N*^*Unlabeled*^ is the number of unlabeled spots in each batch. *N*^*Unlabeled*^=|*S*^*Unlabeled*^|. 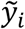 is the pseudo label assigned to each unlabeled spot using the argmax of the model’s prediction, i.e. 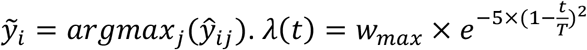 is an epoch-dependent ramp-up function, where *t* is the current epoch, *T* is the ramp-up length, and *w*_*max*_ is the maximum weight. This scheduling strategy allows the model to initially focus on the labeled data and progressively incorporate information from unlabeled data as training proceeds and predictions become more stable.

#### Overall Loss function

GraphScrDom performs representation learning and classification in an end-to-end fashion. The total loss function is designed to jointly minimize reconstruction loss for both gene expression and cell type composition, self-supervised contrastive learning loss, and semi-supervised classification loss. It is defined as:

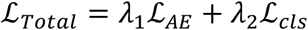

where ℒ_*AE*_ represents the autoencoder loss and ℒ_*cls*_ denotes the classification loss. The autoencoder loss ℒ_*AE*_ incorporates both reconstruction and contrastive learning objectives from the two SCL-GCN autoencoders (for gene expression and cell type composition), as well as the fused representation-driven SCL-GCN reconstruction for gene expression:

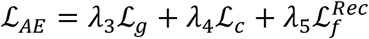

Each modality-specific component ℒ_*g*_ and ℒ_*c*_ includes a reconstruction loss and a contrastive loss:

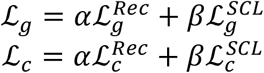

In the above formularization, λ_1_, λ_2_, λ_3_, λ_4_, λ_5_ are hyperparameters controlling the contribution of each loss component, with default values set to 1. The parameters *α* and *β* balance the reconstruction and contrastive learning losses, respectively, and are set by default to *α* = 1 and *β* = 0.1.

### Benchmark methods and evaluation metrics

To evaluate the performance of GraphScrDom, we benchmarked it against seven state-of-the-art methods for spatial domain detection: Leiden, BayesSpace, GraphST, STAGATE, SpaGCN, IRIS, and ScribbleDom. These methods represent a diverse set of algorithms utilizing different data modalities and computational strategies. Leiden, implemented via the SCANPY package (Wolf et al., 2018), is a graph-based community detection algorithm originally designed for single-cell clustering. It has also been widely used for spatial domain identification (Lin et al., 2024). To align the number of clusters with the known number of domains, we employed a resolution search strategy by iteratively adjusting the resolution parameter. BayesSpace is a fully Bayesian model that incorporates a Markov random field prior to account for spatial dependencies, enabling robust spatial clustering (Zhao et al., 2021). GraphST, STAGATE, and SpaGCN are graph-based deep learning models that follow a two-step paradigm: they first learn low-dimensional spot embeddings and then apply unsupervised clustering. SpaGCN integrates gene expression, spatial coordinates, and histology images via a graph convolutional network to construct spot embeddings (Hu et al., 2021). STAGATE utilizes a graph attention autoencoder to learn embeddings by incorporating spatial and transcriptomic information (Dong & Zhang, 2022). GraphST enhances embedding quality with a self-supervised contrastive learning framework built on a graph neural network (Long et al., 2023). IRIS is a multi-modal statistical model that jointly models cell type composition and gene expression to improve spatial domain detection (Ma & Zhou, 2024). ScribbleDom is a weakly supervised model that uses scribble annotations to train a convolutional neural network for spatial domain prediction (Rahman et al., 2023).

To comprehensively assess the performance of all methods, we adopted six commonly used evaluation metrics to quantify the alignment between predicted spatial clusters and ground truth domain labels: Adjusted Rand Index (ARI): measures the similarity between clustering results and true labels, corrected for chance. Normalized Mutual Information (NMI): quantifies the mutual dependence between predicted clusters and true labels. Purity: assesses the extent to which each cluster contains samples from a single ground truth domain. Homogeneity: evaluates whether each predicted cluster contains only samples from a single ground truth class. Completeness: measures whether all samples from a given class are assigned to the same predicted cluster. V-Measure: the harmonic mean of homogeneity and completeness. Together, these metrics offer a robust and multidimensional evaluation of spatial domain detection performance, capturing both accuracy and the structural consistency of predicted domains.

### Software toolkit for scribble generation and spatial domain detection

We have developed GraphScrDom into an easy-to-use and freely accessible software toolkit consisting of two key modules: a scribble annotation tool and a model training tool for spatial domain detection. The scribble annotation tool enables human experts (e.g., pathologists) to manually label spatial domains on spatial grids or histology images via a graphical user interface (GUI) built with PyQt, available as a precompiled executable for macOS and Windows. Users can load a spatial gene expression file and a paired histology image, select regions of interest on the image, and assign domain labels with a few clicks. The tool outputs a scribble-annotated histology image and a corresponding annotated gene expression file, which serve as inputs for model training. The model training tool performs weakly supervised learning using the scribble-annotated data and optionally incorporates a cell type composition file to enable multimodal inference. It outputs a tissue segmentation file with predicted domain labels for all spatial locations. To ensure accessibility, the model training code is available on GitHub (https://github.com/lichen-lab/GraphScrDom), and a simplified version is provided via Google Colab, allowing non-programmers to run the tool with minimal edits. Comprehensive documentation is included on the GitHub page. For instance, applying GraphScrDom to the Human Breast Cancer dataset (3,798 spots and 4 domains) takes approximately 18 minutes per slice with one NVIDIA T4 GPU on Google Colab. The runtime may vary depending on system specifications.

### Simulation strategy

To evaluate GraphScrDom’s ability to integrate both spot-level gene expression and cell type composition for accurate spatial domain detection, we designed a simulation strategy that assesses performance when both modalities are informative and robustness when one modality is degraded or noisy. The simulation is based on the #151507 sample from the Human DLPFC dataset and involves three main steps. (i) Spatial layout and scribble generation: We simulated a spatial layout comprising three distinct domains using the real spatial coordinates from the #151507 sample, and scribble annotations were generated to reflect the ground-truth spatial domains. (ii) Simulated domain-specific gene expression: We used domain-specific spatially variable genes (SVGs) to simulate domain-specific gene expression patterns. Specifically, the top 17 SVGs from each of the three domains in the #151507 sample were selected (51 SVGs in total), and their aggregated expression per spot defined the simulated expression. SVGs were expressed only in their assigned domain, and Gaussian noise was added across all spots to reduce the dominance of domain-specific patterns, with two noise levels: low (σ=3) and high (σ=25), both with mean zero. (iii) Simulated cell type composition: We assumed five cell types per spot and simulated domain-specific compositions using Dirichlet distributions. Domain 1 used Dir(10,5,5,1,1), Domain 2 used Dir(1,5,10,5,1), and Domain 3 used Dir(1,1,5,5,10), resulting in distinct enrichment patterns for each domain. Cell type compositions were sampled per spot from the corresponding domain-specific Dirichlet distribution. To control the influence of composition on domain detection, Gaussian noise was added to the simulated compositions at low (σ=0.3) and high (σ=1) levels, both with mean zero. This simulation framework enables systematic evaluation of GraphScrDom’s robustness and integrative capacity under varying levels of noise across modalities.

## Code availability

The source code for data processing, model architecture and model training; the binary compiled GUI annotation toolkit for scribble annotation and notebook in Google Colab can be found at the github site: https://github.com/lichen-lab/GraphScrDom

## Data availability

The original public data used in this work can be accessed through the following links: Human DLPFC data by 10x Visium available at https://research.libd.org/spatialLIBD/, with Human DLPFC snRNA-seq reference data available at https://app.globus.org/file-manager?origin_id=6f9322c4-5eaf-11ed-b0b5-bfe7e7197080&origin_path=%2F; Human Breast Cancer data by 10x Visium are available at https://www.10xgenomics.com/datasets/human-breast-cancer-block-a-section-1-1-standard-1-0-0, with Human Breast Cancer scRNA-seq reference data at GEO accession GSE176078; Mouse Frontal Cortex data by MERFISH are available at https://cellxgene.cziscience.com/collections/31937775-0602-4e52-a799-b6acdd2bac2e, with snRNA-seq reference data at https://cellxgene.cziscience.com/collections/31937775-0602-4e52-a799-b6acdd2bac2e; Mouse Somatosensory Cortex data by osmFISH are available at https://linnarssonlab.org/osmFISH/availability/, and Mouse Primary Visual Area data by BaristaSeq are available at https://spacetx.github.io/data.html, with same scRNA-seq reference data available at https://cellxgene.cziscience.com/collections/e3aa612b-0d7d-4d3f-bbea-b8972a74dd4b; Mouse Visual Cortex data by STARmap are available at https://clarityresourcecenter.org, with scRNA-seq reference data available at GEO accession GSE102827. Details of the datasets analyzed in this paper are described in Supplement Table 1 and 2.

## Acknowledgement

This study was supported by National Institute of Health (NIH) grants R35GM142701 to L.C. R01AG066653, R01CA266004, R01AG078702, RM1NS133593, 1R01CA288696-01V-Scholar Grant, to R.C.S.

