## Supplementary file for "Reference-Informed Spatial Domain Detection Using Weak Supervision for Spatial Transcriptomics"

### Supplementary Materials for Reference-Informed Spatial Domains Detection Using Weak Supervision for Spatial Transcriptomics

Xin Ma[1,3] , Weijia Jin[1], Qing Lu[1], Ramon C. Sun[2,3 *], Li Chen[1,3 *]

[1] Department of Biostatistics College of Public Health and Health Professions & College of Medicine, University of Florida, Gainesville, FL, USA
[2] Department of Biochemistry & Molecular Biology, College of Medicine, University of Florida, Gainesville, FL, USA
[3] Center for Advanced Spatial Biomolecule Research, University of Florida, Gainesville, FL, USA
* These authors jointly supervised this work: Ramon C. Sun; Li Chen;

### Supplementary Figure

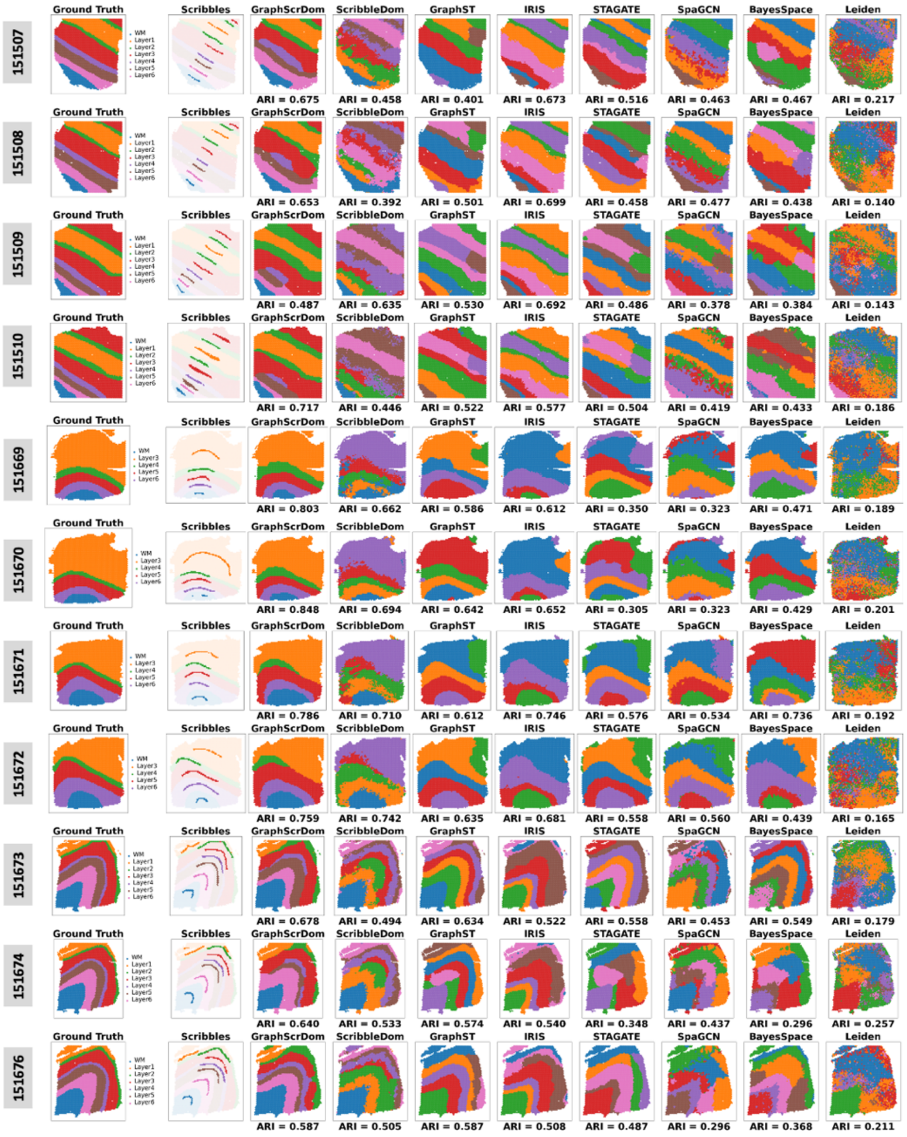

**Figure S1. Evaluation of GraphScrDom on Visium spatial transcriptomics datasets.** Spatial domain detection results on #151507–#151510 for donor 1, #151569–#151572 for donor 2, and #151673, #151674,#151676 for donor 3 from the Human DLPFC dataset. The first two columns display ground truth annotations and scribble inputs. The remaining columns show predicted domains from GraphScrDom, ScribbleDom, GraphST, IRIS, STAGATE, SpaGCN, BayesSpace, and Leiden.

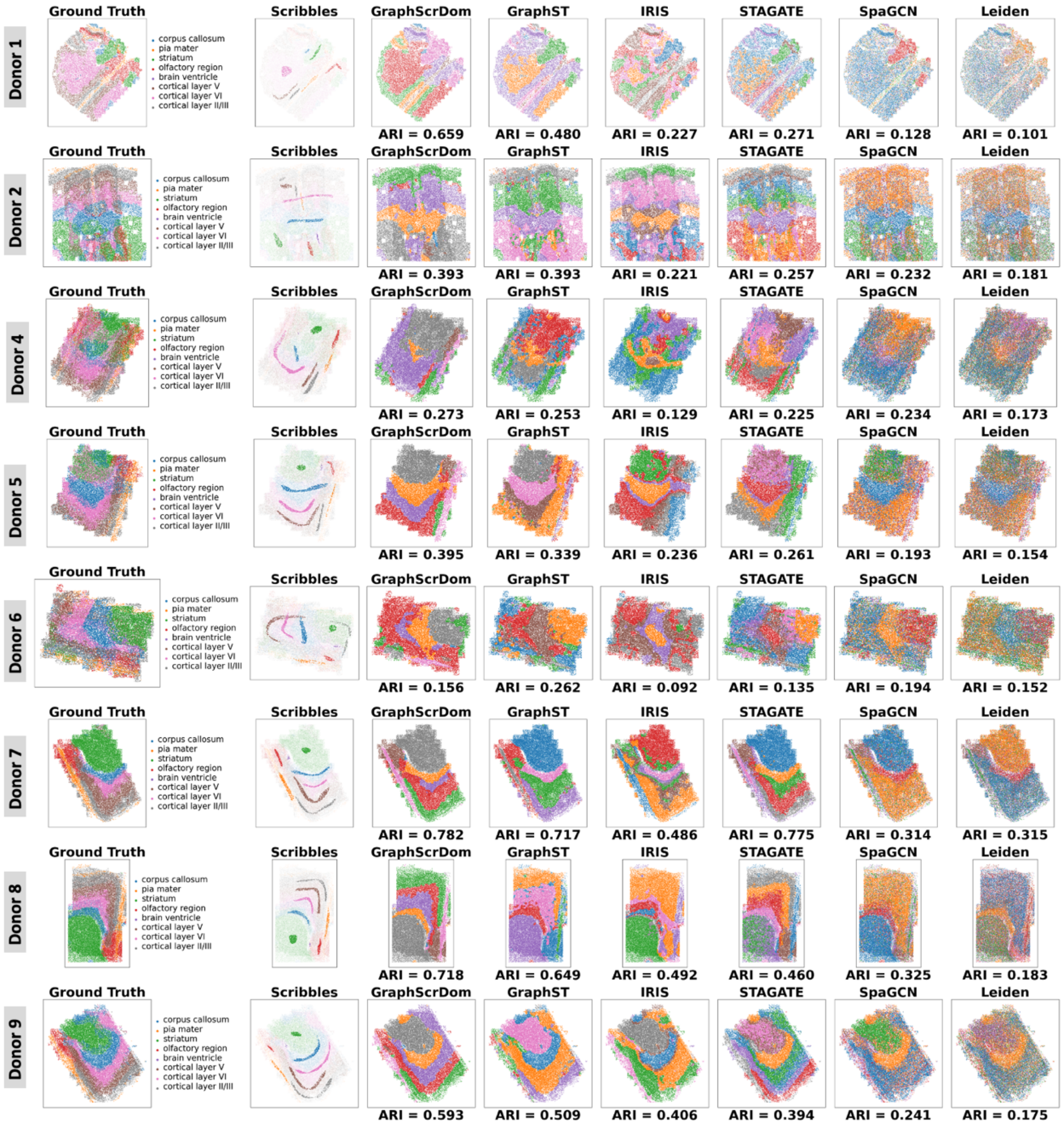

Donor 1
Donor 2
Donor 4
Donor 5
Donor 6
Donor 7
Donor 8
Donor 9
Ground Truth
Scribbles
GraphScrDom
GraphST
IRIS
STAGATE
SpaGCN
Leiden
corpus callosum
pia mater
striatum
olfactory region
brain ventricle
cortical layer V
cortical layer VI
cortical layer II/III
ARI = 0.659
ARI = 0.480
ARI = 0.227
ARI = 0.271
ARI = 0.128
ARI = 0.101
ARI = 0.393
ARI = 0.393
ARI = 0.221
ARI = 0.257
ARI = 0.232
ARI = 0.181
ARI = 0.273
ARI = 0.253
ARI = 0.129
ARI = 0.225
ARI = 0.234
ARI = 0.173
ARI = 0.395
ARI = 0.339
ARI = 0.236
ARI = 0.261
ARI = 0.193
ARI = 0.154
ARI = 0.156
ARI = 0.262
ARI = 0.092
ARI = 0.135
ARI = 0.194
ARI = 0.152
ARI = 0.782
ARI = 0.717
ARI = 0.486
ARI = 0.775
ARI = 0.314
ARI = 0.315
ARI = 0.718
ARI = 0.649
ARI = 0.492
ARI = 0.460
ARI = 0.325
ARI = 0.183
ARI = 0.593
ARI = 0.509
ARI = 0.406
ARI = 0.394
ARI = 0.241
ARI = 0.175

**Figure S2. Evaluation of GraphScrDom on the MERFISH Mouse Frontal Cortex dataset. A.** Spatial domain detection results for eight donors. The first two columns display ground truth annotations and scribble inputs. The remaining columns show predicted spatial domains generated by GraphScrDom and other benchmark methods.

### Supplementary Tables

| Dataset | Sample | Platform | # of Spots/Cells | # of Genes | # of Domains | # of labeled spots (labeled proportion %) | Link |
|---|---|---|---|---|---|---|---|
| Human DLPFC | 151507 | 10x Visium | 4226 | 33538 | 7 | 215 (5.09%) | https://research.libd.org/spatialLIBD/ |
| Human DLPFC | 151508 | 10x Visium | 4384 | 33538 | 7 | 196 (4.47%) | |
| Human DLPFC | 151509 | 10x Visium | 4789 | 33538 | 7 | 231 (4.82%) | |
| Human DLPFC | 151510 | 10x Visium | 4634 | 33538 | 7 | 366 (7.90%) | |
| Human DLPFC | 151669 | 10x Visium | 3661 | 33538 | 5 | 155 (4.23%) | |
| Human DLPFC | 151670 | 10x Visium | 3498 | 33538 | 5 | 220 (6.29%) | |
| Human DLPFC | 151671 | 10x Visium | 4110 | 33538 | 5 | 168 (4.09%) | |
| Human DLPFC | 151672 | 10x Visium | 4015 | 33538 | 5 | 191 (4.76%) | |
| Human DLPFC | 151673 | 10x Visium | 3639 | 33538 | 7 | 291 (8.00%) | |
| Human DLPFC | 151674 | 10x Visium | 3673 | 33538 | 7 | 295 (8.03%) | |
| Human DLPFC | 151675 | 10x Visium | 3592 | 33538 | 7 | 143 (3.98%) | |
| Human DLPFC | 151676 | 10x Visium | 3460 | 33538 | 7 | 257 (7.75%) | |
| Human Breast Cancer | Block A Section 1 | 10x Visium | 3798 | 36601 | 4 | 415 (10.95%) | https://www.10xgenomics.com/datasets/human-breast-cancer-block-a-section-1-1-standard-1-0-0 |
| Mouse Visual Cortex | N/A | STARmap | 1207 | 1020 | 7 | 145 (12.01%) | https://clarityresourcecenter.org |
| Mouse Somatosensory Cortex | N/A | osmFISH | 4839 | 33 | 11 | 555 (11.47%) | https://linnarssonlab.org/osmFISH/availability/ |
| Mouse Primary Visual Area | N/A | BaristaSeq | 4432 | 76 | 6 | 482 (10.88%) | https://spacetx.github.io/data.html |
| Mouse Frontal Cortex | Donor 1 | MERFISH | 21002 | 374 | 8 | 1099 (5.23%) | https://cellxgene.cziscience.com/collections/31937775-0602-4e52-a799-b6acdd2bac2e |
| Mouse Frontal Cortex | Donor 2 | MERFISH | 40484 | 374 | 8 | 1541 (3.81%) | |
| Mouse Frontal Cortex | Donor 3 | MERFISH | 35347 | 374 | 8 | 3012 (8.52%) | |
| Mouse Frontal Cortex | Donor 4 | MERFISH | 31508 | 374 | 8 | 2381 (7.56%) | |
| Mouse Frontal Cortex | Donor 5 | MERFISH | 29371 | 374 | 8 | 2276 (7.75%) | |
| Mouse Frontal Cortex | Donor 6 | MERFISH | 25907 | 374 | 8 | 1643 (6.34%) | |
| Mouse Frontal Cortex | Donor 7 | MERFISH | 32792 | 374 | 8 | 3123 (9.52%) | |
| Mouse Frontal Cortex | Donor 8 | MERFISH | 38338 | 374 | 8 | 2928 (7.64%) | |
| Mouse Frontal Cortex | Donor 9 | MERFISH | 33868 | 374 | 8 | 2887 (8.52%) | |

**Supplementary Table S1. Summary of spatial transcriptomics datasets**

| ST Dataset | Sample | Ref Dataset Type | # of cells/nuclei | # of genes | # cell types | link |
|---|---|---|---|---|---|---|
| Human DLPFC | N/A | snRNAseq | 77581 | 25321 | 25 | https://app.globus.org/file-manager?origin_id=6f9322c4-5eaf-11ed-b0b5-bfe7e7197080&origin_path=%2F |
| Human Breast Cancer | N/A | scRNASeq | 100064 | 29067 | 18 | https://www.ncbi.nlm.nih.gov/geo/query/acc.cgi?acc=GSE176078 |
| Mouse Visual Cortex | N/A | scRNASeq | 2219 | 25202 | 31 | https://www.ncbi.nlm.nih.gov/geo/query/acc.cgi?acc=GSE102827 |
| Mouse Somatosensory Cortex | N/A | scRNASeq | 7531 | 35433 | 21 | https://cellxgene.cziscience.com/collections/e3aa612b-0d7d-4d3f-bbea-b8972a74dd4b |
| Mouse Primary Visual Area | N/A | scRNASeq | 15572 | 35433 | 23 | https://cellxgene.cziscience.com/collections/e3aa612b-0d7d-4d3f-bbea-b8972a74dd4b |
| Mouse Frontal Cortex | Donor 1 | snRNAseq | 21728 | 20984 | 13 | https://cellxgene.cziscience.com/collections/31937775-0602-4e52-a799-b6acdd2bac2e |
| Mouse Frontal Cortex | Donor 2 | snRNAseq | 20152 | 20984 | 13 | |
| Mouse Frontal Cortex | Donor 3 | snRNAseq | 20152 | 20984 | 13 | |
| Mouse Frontal Cortex | Donor 4 | snRNAseq | 21728 | 20984 | 13 | |
| Mouse Frontal Cortex | Donor 5 | snRNAseq | 20152 | 20984 | 13 | |
| Mouse Frontal Cortex | Donor 6 | snRNAseq | 20152 | 20984 | 13 | |
| Mouse Frontal Cortex | Donor 7 | snRNAseq | 21728 | 20984 | 13 | |
| Mouse Frontal Cortex | Donor 8 | snRNAseq | 21728 | 20984 | 13 | |
| Mouse Frontal Cortex | Donor 9 | snRNAseq | 20152 | 20984 | 13 | |

**Supplementary Table S2. Summary of sc(sn)RNAseq reference datasets**

| Platform | Dataset | Sample | GraphScrDom | ScribbleDom | GraphST | IRIS | STAGATE | SpaGCN | BayesSpace | Leiden |
|---|---|---|---|---|---|---|---|---|---|---|
| Visium | Human DLPFC | 151507 | **0.675** | 0.458 | 0.401 | 0.673 | 0.516 | 0.485 | 0.467 | 0.217 |
| Visium | Human DLPFC | 151508 | 0.653 | 0.392 | 0.501 | **0.699** | 0.458 | 0.503 | 0.438 | 0.140 |
| Visium | Human DLPFC | 151509 | 0.487 | 0.635 | 0.530 | **0.692** | 0.486 | 0.392 | 0.384 | 0.143 |
| Visium | Human DLPFC | 151510 | **0.717** | 0.446 | 0.522 | 0.577 | 0.504 | 0.433 | 0.433 | 0.186 |
| Visium | Human DLPFC | 151669 | **0.803** | 0.662 | 0.586 | 0.612 | 0.350 | 0.337 | 0.471 | 0.189 |
| Visium | Human DLPFC | 151670 | **0.847** | 0.694 | 0.642 | 0.652 | 0.305 | 0.332 | 0.429 | 0.201 |
| Visium | Human DLPFC | 151671 | **0.785** | 0.710 | 0.612 | 0.746 | 0.576 | 0.558 | 0.736 | 0.192 |
| Visium | Human DLPFC | 151672 | **0.759** | 0.742 | 0.635 | 0.681 | 0.558 | 0.584 | 0.439 | 0.165 |
| Visium | Human DLPFC | 151673 | **0.678** | 0.494 | 0.634 | 0.522 | 0.558 | 0.470 | 0.549 | 0.179 |
| Visium | Human DLPFC | 151674 | **0.640** | 0.533 | 0.574 | 0.540 | 0.348 | 0.454 | 0.296 | 0.257 |
| Visium | Human DLPFC | 151675 | **0.669** | 0.519 | 0.530 | 0.521 | 0.383 | 0.473 | 0.530 | 0.235 |
| Visium | Human DLPFC | 151676 | **0.587** | 0.505 | **0.587** | 0.508 | 0.487 | 0.306 | 0.368 | 0.211 |
| Visium | Human Breast Cancer | N/A | **0.554** | 0.202 | 0.012 | 0.201 | 0.014 | 0.172 | 0.015 | -0.001 |
| MERFISH | Mouse Frontal Cortex | Donor 1 | **0.659** | N/A | 0.480 | 0.227 | 0.271 | 0.188 | N/A | 0.101 |
| MERFISH | Mouse Frontal Cortex | Donor 2 | **0.393** | N/A | **0.393** | 0.221 | 0.257 | 0.275 | N/A | 0.181 |
| MERFISH | Mouse Frontal Cortex | Donor 3 | **0.674** | N/A | 0.567 | 0.443 | 0.488 | 0.343 | N/A | 0.272 |
| MERFISH | Mouse Frontal Cortex | Donor 4 | **0.273** | N/A | 0.253 | 0.129 | 0.225 | 0.255 | N/A | 0.173 |
| MERFISH | Mouse Frontal Cortex | Donor 5 | **0.395** | N/A | 0.339 | 0.236 | 0.261 | 0.225 | N/A | 0.154 |
| MERFISH | Mouse Frontal Cortex | Donor 6 | 0.156 | N/A | **0.262** | 0.092 | 0.135 | 0.197 | N/A | 0.152 |
| MERFISH | Mouse Frontal Cortex | Donor 7 | **0.782** | N/A | 0.717 | 0.486 | 0.775 | 0.460 | N/A | 0.315 |
| MERFISH | Mouse Frontal Cortex | Donor 8 | **0.718** | N/A | 0.649 | 0.492 | 0.460 | 0.448 | N/A | 0.183 |
| MERFISH | Mouse Frontal Cortex | Donor 9 | **0.593** | N/A | 0.509 | 0.406 | 0.394 | 0.316 | N/A | 0.175 |
| osmFISH | Mouse Somatosensory Cortex | N/A | **0.721** | N/A | 0.512 | 0.180 | 0.472 | 0.437 | N/A | 0.407 |
| BaristaSeq | Mouse Primary Visual Area | N/A | **0.810** | N/A | 0.398 | 0.249 | 0.706 | 0.326 | N/A | 0.198 |
| STARmap | Mouse Visual Cortex | N/A | **0.761** | N/A | 0.557 | 0.465 | 0.651 | 0.440 | N/A | 0.302 |

**Supplementary Table S3. ARI of benchmark methods across all spatial transcriptomics datasets**

| Platform | Dataset | Sample | GraphScrDom | ScribbleDom | GraphST | IRIS | STAGATE | SpaGCN | BayesSpace | Leiden |
|---|---|---|---|---|---|---|---|---|---|---|
| Visium | Human DLPFC | 151507 | **0.734** | 0.596 | 0.622 | 0.732 | 0.693 | 0.614 | 0.626 | 0.319 |
| Visium | Human DLPFC | 151508 | **0.705** | 0.556 | 0.647 | 0.746 | 0.622 | 0.623 | 0.600 | 0.231 |
| Visium | Human DLPFC | 151509 | **0.640** | 0.667 | 0.683 | 0.709 | 0.662 | 0.598 | 0.594 | 0.259 |
| Visium | Human DLPFC | 151510 | **0.731** | 0.585 | 0.649 | 0.647 | 0.613 | 0.591 | 0.562 | 0.281 |
| Visium | Human DLPFC | 151669 | **0.753** | 0.574 | 0.647 | 0.589 | 0.565 | 0.541 | 0.610 | 0.163 |
| Visium | Human DLPFC | 151670 | **0.760** | 0.619 | 0.655 | 0.617 | 0.506 | 0.482 | 0.555 | 0.201 |
| Visium | Human DLPFC | 151671 | **0.745** | 0.609 | 0.719 | 0.708 | 0.704 | 0.667 | 0.692 | 0.210 |
| Visium | Human DLPFC | 151672 | **0.737** | 0.680 | 0.721 | 0.690 | 0.675 | 0.686 | 0.596 | 0.132 |
| Visium | Human DLPFC | 151673 | 0.722 | 0.638 | **0.732** | 0.641 | 0.686 | 0.592 | 0.688 | 0.308 |
| Visium | Human DLPFC | 151674 | **0.713** | 0.647 | 0.702 | 0.661 | 0.460 | 0.549 | 0.482 | 0.307 |
| Visium | Human DLPFC | 151675 | **0.734** | 0.615 | 0.627 | 0.643 | 0.553 | 0.597 | 0.685 | 0.280 |
| Visium | Human DLPFC | 151676 | **0.696** | 0.617 | 0.680 | 0.625 | 0.636 | 0.472 | 0.567 | 0.297 |
| Visium | Human Breast Cancer | N/A | **0.510** | 0.326 | 0.222 | 0.306 | 0.230 | 0.290 | 0.187 | 0.221 |
| MERFISH | Mouse Frontal Cortex | Donor 1 | **0.636** | N/A | 0.587 | 0.364 | 0.455 | 0.234 | N/A | 0.125 |
| MERFISH | Mouse Frontal Cortex | Donor 2 | **0.417** | N/A | 0.380 | 0.327 | 0.340 | 0.256 | N/A | 0.212 |
| MERFISH | Mouse Frontal Cortex | Donor 3 | **0.674** | N/A | 0.620 | 0.517 | 0.605 | 0.361 | N/A | 0.295 |
| MERFISH | Mouse Frontal Cortex | Donor 4 | 0.294 | N/A | 0.244 | 0.209 | **0.325** | 0.221 | N/A | 0.242 |
| MERFISH | Mouse Frontal Cortex | Donor 5 | **0.459** | N/A | 0.395 | 0.321 | 0.374 | 0.230 | N/A | 0.186 |
| MERFISH | Mouse Frontal Cortex | Donor 6 | 0.263 | N/A | **0.275** | 0.180 | 0.229 | 0.198 | N/A | 0.205 |
| MERFISH | Mouse Frontal Cortex | Donor 7 | **0.692** | N/A | 0.652 | 0.475 | 0.674 | 0.371 | N/A | 0.280 |
| MERFISH | Mouse Frontal Cortex | Donor 8 | **0.650** | N/A | 0.589 | 0.497 | 0.552 | 0.352 | N/A | 0.270 |
| MERFISH | Mouse Frontal Cortex | Donor 9 | **0.616** | N/A | 0.537 | 0.467 | 0.510 | 0.315 | N/A | 0.240 |
| osmFISH | Mouse Somatosensory Cortex | N/A | **0.737** | N/A | 0.638 | 0.396 | 0.663 | 0.506 | N/A | 0.447 |
| BaristaSeq | Mouse Primary Visual Area | N/A | **0.811** | N/A | 0.527 | 0.398 | 0.750 | 0.332 | N/A | 0.196 |
| STARmap | Mouse Visual Cortex | N/A | **0.785** | N/A | 0.670 | 0.625 | 0.709 | 0.478 | N/A | 0.326 |

**Supplementary Table S4. NMI of benchmark methods across all spatial transcriptomics datasets**

| Platform | Dataset | Sample | GraphScrDom | ScribbleDom | GraphST | IRIS | STAGATE | SpaGCN | BayesSpace | Leiden |
|---|---|---|---|---|---|---|---|---|---|---|
| Visium | Human DLPFC | 151507 | **0.819** | 0.711 | 0.630 | 0.807 | 0.720 | 0.695 | 0.659 | 0.496 |
| Visium | Human DLPFC | 151508 | 0.801 | 0.633 | 0.731 | **0.811** | 0.697 | 0.725 | 0.709 | 0.440 |
| Visium | Human DLPFC | 151509 | 0.737 | **0.802** | 0.758 | 0.800 | 0.741 | 0.733 | 0.692 | 0.543 |
| Visium | Human DLPFC | 151510 | **0.845** | 0.711 | 0.755 | 0.730 | 0.756 | 0.717 | 0.687 | 0.549 |
| Visium | Human DLPFC | 151669 | **0.878** | 0.779 | 0.817 | 0.780 | 0.803 | 0.777 | 0.818 | 0.632 |
| Visium | Human DLPFC | 151670 | **0.901** | 0.838 | 0.845 | 0.836 | 0.832 | 0.785 | 0.832 | 0.688 |
| Visium | Human DLPFC | 151671 | 0.864 | 0.809 | **0.890** | 0.846 | 0.880 | 0.879 | 0.842 | 0.550 |
| Visium | Human DLPFC | 151672 | 0.850 | 0.852 | **0.882** | 0.801 | 0.851 | 0.872 | 0.710 | 0.473 |
| Visium | Human DLPFC | 151673 | 0.821 | 0.711 | **0.832** | 0.740 | 0.764 | 0.701 | 0.779 | 0.479 |
| Visium | Human DLPFC | 151674 | 0.793 | 0.733 | **0.795** | 0.746 | 0.614 | 0.697 | 0.575 | 0.494 |
| Visium | Human DLPFC | 151675 | **0.805** | 0.706 | 0.694 | 0.722 | 0.572 | 0.706 | 0.721 | 0.448 |
| Visium | Human DLPFC | 151676 | 0.759 | 0.718 | **0.762** | 0.733 | 0.700 | 0.569 | 0.603 | 0.460 |
| Visium | Human Breast Cancer | N/A | **0.803** | 0.572 | 0.511 | 0.595 | 0.517 | 0.555 | 0.477 | 0.511 |
| MERFISH | Mouse Frontal Cortex | Donor 1 | **0.808** | N/A | 0.711 | 0.597 | 0.648 | 0.475 | N/A | 0.427 |
| MERFISH | Mouse Frontal Cortex | Donor 2 | **0.627** | N/A | 0.611 | 0.562 | 0.590 | 0.470 | N/A | 0.443 |
| MERFISH | Mouse Frontal Cortex | Donor 3 | **0.818** | N/A | 0.751 | 0.670 | 0.765 | 0.531 | N/A | 0.527 |
| MERFISH | Mouse Frontal Cortex | Donor 4 | **0.571** | N/A | 0.525 | 0.458 | 0.563 | 0.480 | N/A | 0.476 |
| MERFISH | Mouse Frontal Cortex | Donor 5 | **0.672** | N/A | 0.608 | 0.549 | 0.585 | 0.475 | N/A | 0.449 |
| MERFISH | Mouse Frontal Cortex | Donor 6 | 0.469 | N/A | **0.519** | 0.381 | 0.485 | 0.426 | N/A | 0.422 |
| MERFISH | Mouse Frontal Cortex | Donor 7 | **0.844** | N/A | 0.749 | 0.640 | 0.838 | 0.573 | N/A | 0.498 |
| MERFISH | Mouse Frontal Cortex | Donor 8 | **0.795** | N/A | 0.698 | 0.620 | 0.766 | 0.532 | N/A | 0.487 |
| MERFISH | Mouse Frontal Cortex | Donor 9 | **0.778** | N/A | 0.682 | 0.623 | 0.724 | 0.513 | N/A | 0.461 |
| osmFISH | Mouse Somatosensory Cortex | N/A | **0.834** | N/A | 0.706 | 0.483 | 0.765 | 0.661 | N/A | 0.612 |
| BaristaSeq | Mouse Primary Visual Area | N/A | **0.917** | N/A | 0.530 | 0.589 | 0.886 | 0.620 | N/A | 0.479 |
| STARmap | Mouse Visual Cortex | N/A | **0.877** | N/A | 0.761 | 0.705 | 0.814 | 0.659 | N/A | 0.551 |

**Supplementary Table S5. Purity of benchmark methods across all spatial transcriptomics datasets**

| Platform | Dataset | Sample | GraphScrDom | ScribbleDom | GraphST | IRIS | STAGATE | SpaGCN | BayesSpace | Leiden |
|---|---|---|---|---|---|---|---|---|---|---|
| Visium | Human DLPFC | 151507 | 0.729 | 0.583 | 0.672 | **0.761** | 0.695 | 0.634 | 0.646 | 0.327 |
| Visium | Human DLPFC | 151508 | 0.704 | 0.545 | 0.650 | **0.765** | 0.607 | 0.628 | 0.602 | 0.240 |
| Visium | Human DLPFC | 151509 | 0.649 | 0.659 | 0.662 | **0.720** | 0.632 | 0.565 | 0.569 | 0.265 |
| Visium | Human DLPFC | 151510 | **0.753** | 0.588 | 0.653 | 0.653 | 0.587 | 0.579 | 0.554 | 0.272 |
| Visium | Human DLPFC | 151669 | **0.739** | 0.546 | 0.604 | 0.622 | 0.500 | 0.489 | 0.562 | 0.164 |
| Visium | Human DLPFC | 151670 | **0.736** | 0.586 | 0.627 | 0.633 | 0.439 | 0.428 | 0.496 | 0.175 |
| Visium | Human DLPFC | 151671 | **0.748** | 0.598 | 0.687 | 0.742 | 0.680 | 0.631 | 0.693 | 0.205 |
| Visium | Human DLPFC | 151672 | **0.748** | 0.686 | 0.704 | 0.744 | 0.650 | 0.662 | 0.599 | 0.129 |
| Visium | Human DLPFC | 151673 | **0.725** | 0.620 | 0.716 | 0.674 | 0.677 | 0.586 | 0.674 | 0.316 |
| Visium | Human DLPFC | 151674 | **0.732** | 0.646 | 0.691 | 0.687 | 0.448 | 0.537 | 0.474 | 0.309 |
| Visium | Human DLPFC | 151675 | **0.730** | 0.618 | 0.653 | 0.663 | 0.555 | 0.597 | 0.697 | 0.290 |
| Visium | Human DLPFC | 151676 | **0.690** | 0.618 | 0.674 | 0.628 | 0.635 | 0.464 | 0.567 | 0.314 |
| Visium | Human Breast Cancer | N/A | **0.516** | 0.332 | 0.250 | 0.316 | 0.260 | 0.288 | 0.198 | 0.260 |
| MERFISH | Mouse Frontal Cortex | Donor 1 | **0.657** | N/A | 0.628 | 0.342 | 0.457 | 0.243 | N/A | 0.123 |
| MERFISH | Mouse Frontal Cortex | Donor 2 | **0.456** | N/A | 0.403 | 0.316 | 0.327 | 0.265 | N/A | 0.212 |
| MERFISH | Mouse Frontal Cortex | Donor 3 | **0.683** | N/A | 0.656 | 0.497 | 0.580 | 0.381 | N/A | 0.289 |
| MERFISH | Mouse Frontal Cortex | Donor 4 | **0.337** | N/A | 0.248 | 0.223 | 0.305 | 0.217 | N/A | 0.230 |
| MERFISH | Mouse Frontal Cortex | Donor 5 | **0.470** | N/A | 0.415 | 0.315 | 0.356 | 0.234 | N/A | 0.185 |
| MERFISH | Mouse Frontal Cortex | Donor 6 | 0.290 | N/A | **0.291** | 0.196 | 0.226 | 0.203 | N/A | 0.214 |
| MERFISH | Mouse Frontal Cortex | Donor 7 | **0.702** | N/A | 0.684 | 0.467 | 0.659 | 0.370 | N/A | 0.269 |
| MERFISH | Mouse Frontal Cortex | Donor 8 | **0.653** | N/A | 0.616 | 0.521 | 0.516 | 0.352 | N/A | 0.254 |
| MERFISH | Mouse Frontal Cortex | Donor 9 | **0.641** | N/A | 0.579 | 0.485 | 0.481 | 0.323 | N/A | 0.231 |
| osmFISH | Mouse Somatosensory Cortex | N/A | **0.781** | N/A | 0.668 | 0.391 | 0.632 | 0.497 | N/A | 0.439 |
| BaristaSeq | Mouse Primary Visual Area | N/A | **0.821** | N/A | 0.658 | 0.376 | 0.724 | 0.319 | N/A | 0.195 |
| STARmap | mouse_visual_cortex | N/A | **0.799** | N/A | 0.683 | 0.616 | 0.724 | 0.473 | N/A | 0.321 |

**Supplementary Table S6. Completeness of benchmark methods across all spatial transcriptomics datasets**

| Platform | Dataset | Sample | GraphScrDom | ScribbleDom | GraphST | IRIS | STAGATE | SpaGCN | BayesSpace | Leiden |
|---|---|---|---|---|---|---|---|---|---|---|
| Visium | Human DLPFC | 151507 | **0.740** | 0.609 | 0.579 | 0.704 | 0.692 | 0.596 | 0.608 | 0.312 |
| Visium | Human DLPFC | 151508 | 0.707 | 0.568 | 0.645 | **0.729** | 0.636 | 0.619 | 0.598 | 0.222 |
| Visium | Human DLPFC | 151509 | 0.631 | 0.675 | **0.706** | 0.698 | 0.695 | 0.635 | 0.621 | 0.255 |
| Visium | Human DLPFC | 151510 | **0.710** | 0.582 | 0.646 | 0.641 | 0.640 | 0.604 | 0.571 | 0.291 |
| Visium | Human DLPFC | 151669 | **0.768** | 0.606 | 0.696 | 0.559 | 0.648 | 0.606 | 0.669 | 0.163 |
| Visium | Human DLPFC | 151670 | **0.784** | 0.656 | 0.686 | 0.601 | 0.598 | 0.552 | 0.631 | 0.235 |
| Visium | Human DLPFC | 151671 | 0.742 | 0.621 | **0.755** | 0.678 | 0.730 | 0.707 | 0.691 | 0.216 |
| Visium | Human DLPFC | 151672 | 0.727 | 0.674 | **0.738** | 0.643 | 0.701 | 0.712 | 0.594 | 0.135 |
| Visium | Human DLPFC | 151673 | 0.720 | 0.657 | **0.749** | 0.611 | 0.695 | 0.599 | 0.702 | 0.301 |
| Visium | Human DLPFC | 151674 | 0.695 | 0.647 | **0.713** | 0.637 | 0.473 | 0.562 | 0.490 | 0.305 |
| Visium | Human DLPFC | 151675 | **0.739** | 0.613 | 0.604 | 0.625 | 0.551 | 0.597 | 0.672 | 0.271 |
| Visium | Human DLPFC | 151676 | **0.703** | 0.616 | 0.685 | 0.622 | 0.637 | 0.480 | 0.567 | 0.282 |
| Visium | Human Breast Cancer | N/A | **0.504** | 0.321 | 0.200 | 0.297 | 0.206 | 0.293 | 0.178 | 0.193 |
| MERFISH | Mouse Frontal Cortex | Donor 1 | **0.616** | N/A | 0.550 | 0.389 | 0.453 | 0.226 | N/A | 0.127 |
| MERFISH | Mouse Frontal Cortex | Donor 2 | **0.384** | N/A | 0.359 | 0.339 | 0.353 | 0.248 | N/A | 0.212 |
| MERFISH | Mouse Frontal Cortex | Donor 3 | **0.665** | N/A | 0.587 | 0.540 | 0.633 | 0.343 | N/A | 0.301 |
| MERFISH | Mouse Frontal Cortex | Donor 4 | 0.261 | N/A | 0.241 | 0.196 | **0.347** | 0.225 | N/A | 0.255 |
| MERFISH | Mouse Frontal Cortex | Donor 5 | **0.449** | N/A | 0.377 | 0.328 | 0.395 | 0.227 | N/A | 0.186 |
| MERFISH | Mouse Frontal Cortex | Donor 6 | 0.241 | N/A | **0.260** | 0.167 | 0.232 | 0.193 | N/A | 0.197 |
| MERFISH | Mouse Frontal Cortex | Donor 7 | 0.681 | N/A | 0.624 | 0.483 | **0.689** | 0.372 | N/A | 0.291 |
| MERFISH | Mouse Frontal Cortex | Donor 8 | **0.647** | N/A | 0.565 | 0.475 | 0.594 | 0.352 | N/A | 0.288 |
| MERFISH | Mouse Frontal Cortex | Donor 9 | **0.593** | N/A | 0.500 | 0.451 | 0.543 | 0.307 | N/A | 0.250 |
| osmFISH | Mouse Somatosensory Cortex | N/A | **0.698** | N/A | 0.611 | 0.400 | **0.698** | 0.516 | N/A | 0.455 |
| BaristaSeq | Mouse Primary Visual Area | N/A | **0.801** | N/A | 0.439 | 0.423 | 0.778 | 0.346 | N/A | 0.197 |
| STARmap | Mouse Visual Cortex | N/A | **0.771** | N/A | 0.658 | 0.633 | 0.694 | 0.483 | N/A | 0.331 |

**Supplementary Table S7. Homogeneity of benchmark methods across all spatial transcriptomics datasets**

| Platform | Dataset | Sample | GraphScrDom | ScribbleDom | GraphST | IRIS | STAGATE | SpaGCN | BayesSpace | Leiden |
|---|---|---|---|---|---|---|---|---|---|---|
| Visium | Human DLPFC | 151507 | **0.734** | 0.596 | 0.622 | 0.732 | 0.693 | 0.614 | 0.626 | 0.319 |
| Visium | Human DLPFC | 151508 | 0.705 | 0.556 | 0.647 | **0.746** | 0.622 | 0.623 | 0.600 | 0.231 |
| Visium | Human DLPFC | 151509 | 0.640 | 0.667 | 0.683 | **0.709** | 0.662 | 0.598 | 0.594 | 0.259 |
| Visium | Human DLPFC | 151510 | **0.731** | 0.585 | 0.649 | 0.647 | 0.613 | 0.591 | 0.562 | 0.281 |
| Visium | Human DLPFC | 151669 | **0.753** | 0.574 | 0.647 | 0.589 | 0.565 | 0.541 | 0.610 | 0.163 |
| Visium | Human DLPFC | 151670 | **0.760** | 0.619 | 0.655 | 0.617 | 0.506 | 0.482 | 0.555 | 0.201 |
| Visium | Human DLPFC | 151671 | **0.745** | 0.609 | 0.719 | 0.708 | 0.704 | 0.667 | 0.692 | 0.210 |
| Visium | Human DLPFC | 151672 | **0.737** | 0.680 | 0.721 | 0.690 | 0.675 | 0.686 | 0.596 | 0.132 |
| Visium | Human DLPFC | 151673 | 0.722 | 0.638 | **0.732** | 0.641 | 0.686 | 0.592 | 0.688 | 0.308 |
| Visium | Human DLPFC | 151674 | **0.713** | 0.647 | 0.702 | 0.661 | 0.460 | 0.549 | 0.482 | 0.307 |
| Visium | Human DLPFC | 151675 | **0.734** | 0.615 | 0.627 | 0.643 | 0.553 | 0.597 | 0.685 | 0.280 |
| Visium | Human DLPFC | 151676 | **0.696** | 0.617 | 0.680 | 0.625 | 0.636 | 0.472 | 0.567 | 0.297 |
| Visium | Human Breast Cancer | N/A | **0.510** | 0.326 | 0.222 | 0.306 | 0.230 | 0.290 | 0.187 | 0.221 |
| MERFISH | Mouse Frontal Cortex | Donor 1 | **0.636** | N/A | 0.587 | 0.364 | 0.455 | 0.234 | N/A | 0.125 |
| MERFISH | Mouse Frontal Cortex | Donor 2 | **0.417** | N/A | 0.380 | 0.327 | 0.340 | 0.256 | N/A | 0.212 |
| MERFISH | Mouse Frontal Cortex | Donor 3 | **0.674** | N/A | 0.620 | 0.517 | 0.605 | 0.361 | N/A | 0.295 |
| MERFISH | Mouse Frontal Cortex | Donor 4 | 0.294 | N/A | 0.244 | 0.209 | **0.325** | 0.221 | N/A | 0.242 |
| MERFISH | Mouse Frontal Cortex | Donor 5 | **0.459** | N/A | 0.395 | 0.321 | 0.374 | 0.230 | N/A | 0.186 |
| MERFISH | Mouse Frontal Cortex | Donor 6 | 0.263 | N/A | **0.275** | 0.180 | 0.229 | 0.198 | N/A | 0.205 |
| MERFISH | Mouse Frontal Cortex | Donor 7 | **0.692** | N/A | 0.652 | 0.475 | 0.674 | 0.371 | N/A | 0.280 |
| MERFISH | Mouse Frontal Cortex | Donor 8 | **0.650** | N/A | 0.589 | 0.497 | 0.552 | 0.352 | N/A | 0.270 |
| MERFISH | Mouse Frontal Cortex | Donor 9 | **0.616** | N/A | 0.537 | 0.467 | 0.510 | 0.315 | N/A | 0.240 |
| osmFISH | Mouse Somatosensory Cortex | N/A | **0.737** | N/A | 0.638 | 0.396 | 0.663 | 0.506 | N/A | 0.447 |
| BaristaSeq | Mouse Primary Visual Area | N/A | **0.811** | N/A | 0.527 | 0.398 | 0.750 | 0.332 | N/A | 0.196 |
| STARmap | Mouse Visual Cortex | N/A | **0.785** | N/A | 0.670 | 0.625 | 0.709 | 0.478 | N/A | 0.326 |

**Supplementary Table S8. V-Measure of benchmark methods across all spatial transcriptomics datasets**

| | Median | Avg. Improvement | Avg. Percent Improvement (%) | Win Count | P-Value |
|---|---|---|---|---|---|
| **GraphScrDom** | **0.678** | N/A | N/A | N/A | N/A |
| **ScribbleDom** | 0.526 | 0.126 | 25.84 | 11/12 | 0.005394 |
| **GraphST** | 0.580 | 0.129 | 24.10 | 10/12 | 0.002678 |
| **IRIS** | 0.632 | 0.073 | 13.24 | 10/12 | 0.032630 |
| **STAGATE** | 0.487 | 0.231 | 58.00 | 12/12 | 0.001263 |
| **SpaGCN** | 0.462 | 0.248 | 61.78 | 12/12 | 0.001263 |
| **BayesSpace** | 0.438 | 0.230 | 54.90 | 12/12 | 0.001263 |
| **Leiden** | 0.190 | 0.499 | 267.59 | 12/12 | 0.001263 |

**Supplementary Table S9. Summary statistics (ARI) for all benchmark methods on Human DLPFC Dataset**

| | Median | Avg. Improvement | Avg. Percent Improvement (%) | Win Count | P-Value |
|---|---|---|---|---|---|
| **GraphScrDom** | **0.593** | N/A | N/A | N/A | N/A |
| **ScribbleDom** | NA | NA | NA | N/A | NA |
| **GraphST** | 0.480 | 0.053 | 8.48 | 8/9 | 0.037780 |
| **IRIS** | 0.236 | 0.212 | 80.42 | 9/9 | 0.004576 |
| **STAGATE** | 0.271 | 0.153 | 47.80 | 9/9 | 0.004576 |
| **SpaGCN** | 0.275 | 0.215 | 74.42 | 8/9 | 0.008909 |
| **BayesSpace** | N/A | N/A | N/A | N/A | N/A |
| **Leiden** | 0.175 | 0.326 | 190.37 | 9/9 | 0.004576 |

**Supplementary Table S10. Summary statistics (ARI) for all benchmark methods on Mouse Frontal Cortex Dataset**
